# Circadian regulation of key physiological processes by the RITMO1 clock protein in the marine diatom *Phaeodactylum tricornutum*

**DOI:** 10.1101/2024.12.23.629939

**Authors:** Alessandro Manzotti, Raphaël Monteil, Soizic Cheminant Navarro, Dany Croteau, Lucie Charreton, Antoine Hoguin, Nils Fabian Strumpen, Denis Jallet, Fayza Daboussi, Peter Kroth, François-Yves Bouget, Marianne Jaubert, Benjamin Bailleul, Jean-Pierre Bouly, Angela Falciatore

**Affiliations:** Laboratoire de Biologie du Chloroplaste et Perception de la Lumière chez les Microalgues, UMR7141, CNRS, Sorbonne Université, Institut de Biologie Physico-Chimique, Paris, France; Institute of Molecular Plant Science, University of Edinburgh, Max Born Crescent, Edinburgh EH9 3BF, UK; Fachbereich Biologie, Universität Konstanz, 78457 Konstanz, Germany; Toulouse Biotechnology Institute (TBI), Université de Toulouse, CNRS, INRAE, INSA, Toulouse, France; Toulouse White Biotechnology (TWB), INSA, Toulouse, France; Laboratoire d’Océanographie Microbienne Sorbonne Université, CNRS UMR7621, Observatoire Océanologique de Banyuls, Banyuls sur Mer, France; Molécules de Communication et Adaptation des Micro-Organismes, UMR 7245, CNRS/MNHN, Paris, France

**Keywords:** Diatom, phytoplankton, RITMO1, biological rhythms, circadian clock, photosynthesis, photophysiology, gene expression

## Abstract

- Phasing biological and physiological processes to periodic light-dark cycles is crucial for the life of most organisms. Marine diatoms, as many phytoplanktonic species, exhibit biological rhythms, yet their molecular timekeepers remain largely uncharacterized. Recently, the bHLH-PAS protein RITMO1 has been proposed to act as a regulator of circadian rhythms.
- In this study, we first determined the physiological conditions to monitor circadian clock activity and its perturbation in the diatom model species *Phaeodactylum tricornutum* by using cell fluorescence as a circadian output. Employing ectopic overexpression, targeted gene mutagenesis, and functional complementation, we then investigated the role of RITMO1 in various circadian processes.
- Our findings reveal that RITMO1 significantly influences the *P. tricornutum* circadian rhythms not only of cellular fluorescence, but also of photosynthesis and of the expression of clock-controlled genes, including transcription factors and putative clock input/output components. RITMO1 effects on rhythmicity are unambiguously detectable under free running conditions.
- By uncovering the complex regulation of biological rhythms in *P. tricornutum*, these results provide a key step in understanding the endogenous regulators of phytoplankton physiological responses to environmental changes. Furthermore, these studies position diatoms as instrumental and novel model systems for elucidating key mechanistic principles of oscillator functions in marine ecosystems.

## Introduction

Diatoms are unicellular algae that can colonize all aquatic environments, while also being found in sea ice, mudflats and soils (Pierella Karlusich *et al*., 2020). With an estimated number of more than 100,000 species (Malviya *et al*., 2016), diatoms represent the most species-rich algae, playing a central role in trophic networks and biogeochemical cycles of the oceans. Although being predominantly photosynthetic, diatoms are not directly related to terrestrial plants and green algae of the Archaeplastida clade, which are derived by a primary endosymbiosis event. Like other Stramenopiles, diatoms acquired their plastid through endosymbiotic events involving a eukaryotic red and possibly a green alga taken up by a heterotrophic host cell (Moustafa *et al*., 2009; Burki *et al*., 2020). Traces of this complex evolutionary history are found in their genomes and in the peculiar gene repertoire derived from their heterotrophic and phototrophic ancestors, as well as from bacteria via horizontal gene transfer (Mock *et al*., 2022). Comparative genomics also highlighted diatom core genes and species specific genes, many of unknown function, which may represent genetic innovations of relevance for the ecological success of these microalgae (Bowler et al., 2008; Mock et al., 2022).

Circadian clocks have in common that they constitute timing systems generating biological cycles of ∼24 h. Such clocks have been identified in a wide variety of organisms, from bacteria to humans (Dunlap, 1999; Bell-Pedersen et al., 2005). While these clocks involve many different mechanisms and molecular players, they usually share common features. In eukaryotes, they are mainly structured around interlocked negative transcriptional-translational feed-back loops (TTFL) that use transcription factors to control the expression of cognate genes (Dunlap, 1999). Besides, non-TTFL-based oscillatory systems also exist (Sweeney & Haxo, 1961; Ishiura et al., 1998; Edgar et al., 2012). Periodic light and dark cycles represent the primary time giver (“Zeitgeber time”, ZT), with light-sensing receptors acting in the input pathways to synchronize the circadian clock to diel light rhythms (Millar et al., 1995; Oakenfull & Davis, 2017). In addition to light, other abiotic factors, such as temperature, can play an influential role in clock entrainment (Rensing & Ruoff, 2002). The clock regulatory loop is self-sustained, i.e. oscillations of the endogenous clock, as well as controlled physiological processes, are maintained for several days after exposure to a constant environment with no light/dark (L:D) or temperature cycles. In photosynthetic organisms, the circadian clock system has been extensively characterized in the model plant species *Arabidopsis thaliana*, where intricated regulatory feedback loops control the rhythmicity of major metabolic and cellular processes (Dodd et al., 2005; Fung-Uceda et al., 2018; de Barros Dantas et al., 2023), as well as the developmental transition to flowering (Song et al., 2015). In algae, timekeeper components have so far been elucidated in detail only in representative model species of the green lineage derived by primary endosymbiosis (Petersen et al., 2022), such as the Chlorophyceae *Chlamydomonas reinhardtii* and the Mamiellophyceae *Ostreococcus tauri*. The latter time keeping mechanism relies on a minimalist plant-like clock with only two components, OtCCA1 and OtTOC1 (Corellou et al., 2009).

There is compelling evidence that diatoms, like other phototrophs, have evolved to adjust their physiology and metabolism to daily light/dark fluctuations. Under laboratory conditions, diel rhythms have been described in diverse diatom species for essential processes such as photosynthesis (Post *et al*., 1984; Jallet *et al*., 2016), pigment synthesis (Owens *et al*., 1980; Hunsperger *et al*., 2016), cell division (Nelson & Brand, 1979), and gene expression modulation (Chauton *et al*., 2013; Smith *et al*., 2016; Bilcke *et al*., 2021). *In situ* studies of natural populations also reported rhythms in optical properties related to photosynthetic activity and growth (Brand, 1982; Kheireddine & Antoine, 2014). Accordingly, metatrascriptomic analyses detected diel patterns of gene expression in natural diatom communities (Kolody *et al*., 2019; Coesel *et al*., 2021). Nonetheless, the involvement of endogenous timekeeper(s) in regulation of rhythmicity remains enigmatic in diatoms, mainly because of two methodological difficulties. Firstly, while rhythms have been documented in numerous species under L:D cycles, their persistence under free-running conditions, a hallmark of clock function, has received limited attention (Palmer *et al*., 1964; Nelson & Brand, 1979; Chisholm & Brand, 1981; Ragni & d’Alcalà, 2007; Häfker *et al*., 2023), also due to the lack of appropriate methods for effectively studying circadian processes. Secondly, with the exception of cryptochromes and casein kinases (Coesel *et al*., 2009; Farré, 2020; Jaubert *et al*., 2022), diatom genome sequences lack clear orthologs of key clock genes identified in bacteria, algae, plants or animals. Yet, although highly divergent, several protein domains of eukaryotic clock components can be found in diatom protein databanks. Proteins containing these domains might have acquired molecular functions in circadian clock regulation, as recently proposed for the diatom RITMO1 protein (Annunziata *et al*., 2019), which, similarly to central animal clock proteins (Takahashi, 2017), possesses bHLH-PAS domains. RITMO1 was first identified in the diatom model species *Phaeodactylum tricornutum* in a survey of the most rhythmic transcription factors (Annunziata *et al*., 2019). Since, multiple *RITMO1*-like bHLH-PAS genes have been found in all diatoms and other Stramenopiles for which omic sequences are available, as well as in other phytoplanktonic organisms of different phyla (Annunziata *et al*., 2019; Farré, 2020; Bilcke *et al*., 2021). Deregulation of *RITMO1* expression levels and timing resulted in *P. tricornutum* cells with altered gene expression profiles in L:D cycles but also in continuous darkness (D:D), showing that gene expression rhythmicity affected by *RITMO1* is not directly dependent on light inputs (Annunziata *et al*., 2019). Initial analyses of cellular fluorescence rhythms, which in diatoms reflect synchrony in the plastid ontogeny during cell cycles (Ragni & d’Alcalà, 2007; Huysman *et al*., 2013; Hunsperger *et al*., 2016), also revealed that RITMO1 influences circadian rhythms (Annunziata *et al*., 2019) and suggested a possible function for this protein in the circadian clock. Recently the photoreceptor Aureochrome1a (AUREO1a) of *P. tricornutum*, has been proposed to be involved in diatom circadian regulation. Aureochromes are exclusively found in Stramenopiles (Takahashi *et al*., 2007; Ishikawa *et al*., 2009) and possess both a blue light-sensing LOV domain and a DNA-binding bZIP domain. While AUREO1a is known for its central role in the *P. tricornutum* blue light responses (Kroth *et al*., 2017), PtAUREO1a knock-out lines have been shown to exhibit perturbed rhythmic expression of several genes, including *RITMO1* and Cryptochrome Photolyase *CPF1* in L:D and D:D (Mann *et al*., 2020; Madhuri *et al*., 2024).

In this study, we used RITMO1 mutant lines to further characterize the circadian clock function in diatoms. Alongside *RITMO1* over-expressing lines, we generated RITMO1 loss of function mutants of *P. tricornutum* by CRISPR/Cas9 editing and tested the consequence of these modifications for various biological rhythms under different L:D cycles and free-running conditions. Further supported by complementation tests in *RITMO1* mutants, our findings revealed a tight molecular control of rhythmic events in *P. tricornutum*, highlighting a key role for RITMO1 in the circadian regulation of diatom physiology.

## Materials and Methods

### Cultures and irradiation conditions

Transgenic and control lines of *P. tricornutum* used in this study refer to strain Pt1 8.6; CCMP2561 (Bowler et al., 2008), with the exception of the *PtAUREO1a* knock-out mutants generated in the UTEX646 Pt4 strain by Madhuri et al. (Madhuri *et al*., 2024). Cells were cultured in F/2 media (Guillard, 1975) at 18°C under white light (Bridgelux BXRV-DR-1830H-3000-A-13 LEDs) of different intensities and photoperiods, as described in the text. Cells were preadapted at least two weeks to specific conditions before performing analyses of rhythmicity. For most of the experiments, batch cultures were grown in incubators with orbital agitation and analyses performed in cells in exponential phase of growth (between 0.1-2×10^6^ cell ml^−1^). Temperature compensation experiments were performed on cells entrained under 16L:8D 25 µmol photons m^−2^ s^−1^ at 18°C and subsequently exposed to L:L of 17 µmol photons m^−2^ s-^1^ at different temperature ranges (14°C-22°C). Low temperature shift was done at the last L:D transition (ZT 16) and at the higher temperature at the last D:L transition (ZT 24) (Poliner *et al*., 2019). For studies of photosynthesis, cells were grown in turbidostat modules (Multi-Cultivator MC 1000-OD, Photon Systems Instruments, Czech Republic) and maintained at an optical density corresponding to 1-2×10^6^ cell ml^−1^. For all the experiments, cell concentrations were measured using a Z2 Coulter Particle Count and Size Analyser (Beckman Coulter, USA). Growth capacity of the different strains was calculated during the exponential phase of growth over four days and expressed as number of divisions per day: log2 (C_4_/C_0_)/4, where C_0_ and C_4_ represent the culture concentrations at day 0 and day 4, respectively.

### Generation of RITMO1 mutant strains

RITMO1 mutants were generated by biolistic delivery of the CRISPR-Cas9 ribonucleoproteins according to Serif *et al*., (2018). Selection of transformed lines was accomplished by co-targeting the *APT* gene (Phatr3_J6834) to induce resistance to 2-fluoroadenine (2-FA). Three and two independent sgRNAs were co-transformed to target *RITMO1* and *APT*, respectively (Table S1). Eight 2-FA resistant colonies were sequenced and four of them appeared to bear mutations on the two alleles of *RITMO1*. WT *P. tricornutum* cells, the KO1 and KO2 lines, an additional transgenic line showing no mutation in *RITMO1* (chosen as control, CTR line) have been characterized in this study. Moreover, rhythms have been also studied in a previously generated *RITMO1* ectopic overexpression line (OE1) (Annunziata *et al*., 2019). A RITMO1 complemented line (named KO2-C2) was obtained by transforming KO2 with a construct containing the genomic *RITMO1* sequence fused to the Venus reporter, under the endogenous *RITMO1* promoter and terminator by microparticle bombardment and characterized as described in Method S3.

#### Cellular fluorescence analysis

In circadian experiments, culture samples were automatically collected every 2 h via a BVP standard ISM444 peristaltic pump (Ismatec) controlled by a FC204 fraction collector (Gilson). Prior to cytometry analysis, cells were stored at 4°C in the dark for at least 2 h to a maximum of 24 h. Cellular fluorescence rhythmicity (FL3 parameter) analyses were performed using a MACSQuant flow cytometer (Miltenyi Biotec), (Annunziata et al., 2019). FL3-A parameter (488 nm excitation, 655-730 nm detection) was used to track chlorophyll fluorescence. Between 10,000 and 30,000 events were counted at each time point and for each culture replicate and median FL3-A value were used for following rhythmicity.

### Analysis of photosynthetic parameters

For photophysiology characterisations, cells were grown in turbidostat modules, adapted to 16L:8D (50 µmol photons m^−2^ s^−1^) four days prior to the beginning of sampling and then shifted to continuous light (30 µmol photons m^−2^ s^−1^). Analyses were performed on four replicas of each genotype, measured every fifteen minutes, always in the order WT, KO1, KO2, OE1. Therefore, a given strain was sampled every 4h but each genotype was sampled every hour. The samples remained for 1 min in darkness for relaxation before the measurements in the Fluorescence Induction and Relaxation fluorometer (miniFIRe, Gorbunov *et al*., 2020). A saturating single turnover flash (100 µs) of blue light (450 nm) applied on dark-acclimated samples induced a fluorescence induction from F_0_ (extrapolated minimum fluorescence) to F_m_ (maximum fluorescence), which is then used to calculate the maximum quantum yield as F_v_/F_m_ = (F_m_ - F_0_) / F_m_. Then, a rapid light curve protocol was performed consisting of 9 steps of increasing light intensities (*E*) (from 25 to 800 µmol photons m^−2^ s^−1^) with 1 min exposure duration for the first two light steps, followed by 20 seconds for the 7 next steps. Different photosynthetic parameters were calculated, including the light irradiance at half saturation of the non-photochemical quenching (NPQ) (E50NPQ) and the NPQ relaxation after 4min in the dark (NPQ rel.) as described in the Supplementary Method S1 and Fig. S1.

### Analysis of rhythmic processes

The analysis of FL3-A fluorescence rhythmicity was performed using the BioDare2 webtool (https://biodare2.ed.ac.uk/, Zielinski *et al*., 2014). Data traces were first analysed using the Enright Periodogram (EPR) algorithm, which was used as an arrhythmicity test (Zielinski *et al*., 2014). Tracks not identified as rhythmic by EPR were used for graphical representations of average fluorescence, but were not further analysed for period, amplitude and Relative Amplitude Errors (RAE). Period, amplitude and RAE were calculated via Fast Fourier Transform Non-Linear Least Square Algorithm (FFT-NLLS), within a period range of 18-34 h. At least three full oscillations were considered for the analysis. RAE, defined as the ratio of the amplitude error to the most probable derived amplitude magnitude (Plautz *et al*., 1997), was used as a measure of degree of confidence in the fit used by the algorithm to calculate rhythmic parameters. Statistical differences were examined using unpaired Student’s t-test with the WT as reference sample. For graphical representations, data traces were baseline detrended, normalized to the maximum value and aligned to the mean values.

For the analysis of photosynthesis, due to the constraints of the manual sampling, a limited number of cycles have been analysed, preventing the appropriate use of the BioDare2 tool. Thanks to the high temporal sampling resolution, however, we were able to conduct rhythmicity tests for photosynthesis using the JTK_CYCLE algorithm (v3.1) (Hughes et al., 2010), by using the “replica option”. For the period in 16L:8D conditions, a period of 24 h was enforced. In L:L, because circadian clock free-runs with its own period, which does not match exactly 24 h, the period windows was set between 20 and 28 h.

### RNA extraction and RT-qPCR analysis

For RNA purification, 10^8^ cells from exponential phase cultures were filtered onto Whatman 589/2 filters. RNA was extracted with TriPure Isolation Reagent (Roche) following the provided protocol and further purified by an ammonium acetate precipitation. 500ng of RNA were reverse-transcribed using an oligo-dT and random primers mix and the QuantiTect Reverse Transcription Kit (Qiagen). For RT-qPCR, the 30S ribosomal small subunit protein (*RPS*) and TATA box Binding Protein (*TBP*) were used as reference genes (Siaut *et al*., 2007). For each sample, the geometrical average of Ct was used as reference to evaluate the expression of genes of interest via the ΔΔCt Livak method (Livak & Schmittgen, 2001). For each gene, data was normalized against the maximum expression value among the different strains, so that expression values are all between 0 and 1. This normalization method allows to compare the relative gene expression among different cell lines. RT-qPCR analyses were performed on two biological replicates in L:D samples, and on three replica for experiments in L:L and with the complemented line. Primers for RT-qPCR analysis are listed in Table S1.

## Results

### Unveiling the experimental conditions for characterising the circadian clock in *P. tricornutum*

To define the appropriate experimental conditions to monitor circadian clock activity in *P. tricornutum*, we analysed rhythmicity under different light conditions, by using as circadian output the red fluorescence per cell (FL3-A parameter in the flow cytometer), a proxy of chlorophyll a cellular content (Hunsperger *et al*., 2016; Annunziata *et al*., 2019). First, we analysed rhythmicity in cells adapted to 16L:8D photoperiods under moderate light intensity (25 µmol photons m^−2^s^−1^) resulting in ∼1 cell division per day (Fig 1a, Fig 1g and Table S2). FL3 displayed strong diel oscillations with an acrophase of fluorescence around 13h after light onset (ZT13), hence anticipating by 3h the transition to darkness (Table S2). During the dark period, fluorescence decreased and reached a minimum at the end of the night, when cells are expected to have completed their cell division. The minimum of fluorescence coincided with the peak of cells in G1 phase (Fig. S2 and Annunziata *et al*., 2019), confirming that FL3 is also a proxy for cell division. When grown under 16L:8D cycles with higher light intensity (75 µmol photons m^−2^s^−1^), cells showed strong rhythms (Fig 1b), as for cells experiencing moderate light, with a slightly delayed acrophase of fluorescence (ZT14) and higher growth (Fig 1g and Table S2). L:D cycles with a photoperiod other than 24 h (also called T cycles) are a convenient means for disentangle the effect of external cues and circadian clock on physiological rhythms since the circadian clock is usually not entrained below T=16h (Roenneberg *et al*., 2005). We performed additional experiments by shifting cells adapted to12L:12D at 25 µmol photons m^−2^s^−1^ to 6L:6D cycles (period, T=12h). The analysis in Fig 1c showed the FL3 trend continued to rise during the first 12 hours, as for cells adapted to a 24-hour period. However, a decrease in FL3 was already observed after the first 6h of darkness and, from the second day onwards, FL3 oscillations completely shifted to shorter T12 cycles. Since this coincided with a decrease in growth (0.5 divisions/day) (Fig. 1g and Table S2), it is likely that the observed ultradian fluorescence patterns primarily reflect the inhibitory effect of anticipated dark on the cell cycle. To measure circadian rhythms, we thus focused subsequent analyses in L:D entrained cells exposed to free running conditions, where the contribution of other processes (e.g., cell cycle and direct responses to environmental changes) should be minimized.

**Fig. 1:**
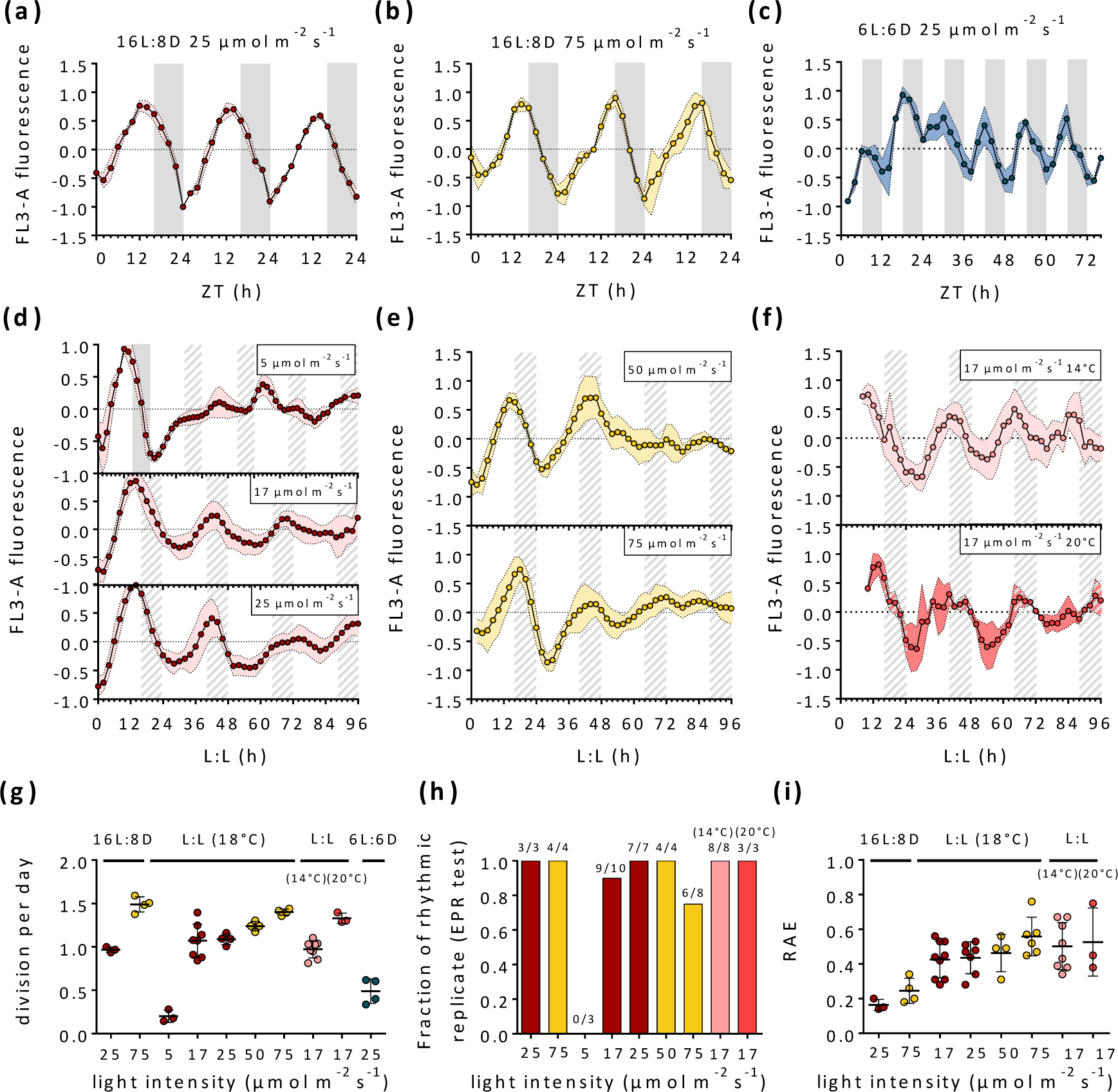
Characterisation of cellular fluorescence rhythmicity of *P. tricornutum* grown under different L:D cycles of different light intensities and periods and following a shift to different free-running conditions. In the figure, red dots represent cells entrained in 16L:8D cycles at 25 µmol photons m⁻² s⁻¹ 18°C, fluorescence rhythms (FL3-A) shown in (a); yellow dots indicate cells entrained under 16L:8D at 75 µmol photons m⁻² s⁻¹ 18°C, with FL3-A in (b). In (c) FL3 trends of cells shifted to 6L:6D cycles at 25 µmol photons m⁻² s⁻¹ 18°C after prior entrainment under 12L:12D at the same intensity. In the middle panels (d-f) free-running FL3 rhythms over four subjective days: d) cells entrained as in (a) and subsequently exposed to L:L of 5 µmol photons m^−2^ s^−1^ (top), 17 µmol photons m^−2^ s^−1^ (middle) and 25 µmol photons m^−2^ (bottom); e) cells grown as in (b) then exposed to L:L at 50 (Top) or to 75 µmol photons m^−2^ s^−1^ (bottom). In (f), cells were entrained as in (a) and then exposed to L:L of 17 µmol photons m^−2^ s^−1^ at 14°C and 20°C. Cell division per day calculated in the different experiments (n>=3) is reported in (g); Statistical analysis of rhythmicity, indicating the fraction of the biological replica (n>=3) found rhythmic by EPR test and so analysed with FFT-NLLS (Fast Fourier Transform Non-Linear Least Square Algorithm) (h) and (i) the RAE (Relative Amplitude of Error). Fluorescence profiles were baseline detrended and then normalized between 1 and −1 for graphic representation. Colored dashed lines represent standard deviations. White and grey regions represent light and dark periods, grey dashed regions represent subjective nights in L:L. ZT indicates the “*Zeitgeber time*” in light–dark regime. In L:L, the time is counted starting from the end of the last L:D day.

We first analyzed responses of cells adapted to moderate 16L:8D as in Fig 1a and then exposed to either continuous dark (D:D) (Table S2) or continuous white light (L:L) of various intensities for four days (Fig 1d). The transition to D:D resulted in total flattening of fluorescence signal after the first subjective night (Table S2), as expected upon cell division arrest in darkness (Huysman *et al*., 2010). When transferred to L:L under low light intensity (5 µmol photons m^−2^ s^−1^, Fig. 1d, top panel), cells continued to divide over the whole four-day period, although at a strongly reduced rate when compared to higher light intensities (Fig. 1g) and the FL3 rhythmicity dampened (Table S2). None of the biological replicates appeared as rhythmic based on our circadian periodicity test (Fig. 1h, Table S2). In the case of cultures exposed to 17 µmol photons m^−2^ s^−1^ L:L (Fig. 1d, middle), and thus experiencing during the 24 h a total irradiance approximately equivalent to that previously experienced in 16L:8D cycle, the oscillations showed a robust rhythmic pattern. They persisted for at least 4 days in L:L, although with a longer period as compared to L:D (around 28 h, Table S2). FL3 oscillations also persisted when the light intensity was maintained at 25 µmol photons.m^−2^.s^−1^ in L:D and L:L (Fig. 1d, bottom, Table S2).

When cells were acclimated to a 16L:8D photoperiod under higher light intensity (75 µmol photons m ^−2^ s^−1^) as in Fig. 1b and then were transferred to L:L of 50 µmol photons m^−2^ s^−1^ (Fig 1e, top and Table S2), oscillations were visible though only during the first two days of free-run, with the amplitude decreasing and the noise in the period calculation increasing subsequently (Fig 1h and i and Table S2). The rhythmic profiles became even more perturbated in cells exposed to a higher light intensity of 75 µmol photons m^−2^ s^−1^in L:L (Fig. 1e, h and i).

A typical circadian clock feature is the ability to maintain rhythms under a wide range of temperatures. Therefore, we also tested FL3 responses in cells adapted to 16L:8D of moderate light at the growth condition (18°C) and then exposed to L:L with temperature ranging from 14 to 22°C (Fig. 1f and Fig. S3). *P. tricornutum* showed robust circadian rhythms at all the temperatures below 18°C, with slightly shorter periods at 14°C and 16°C. Similar rhythms were detectable at 18°C and 20°C, but they were largely affected at higher temperatures (Fig. S3). When detectable, they showed reduced amplitudes.

Taken together, these experiments supported the presence of a circadian oscillator in diatoms and identified the experimental conditions (Fig. 1d) for analysing its activity and consequence of its perturbation in *P. tricornutum RITMO1* mutants.

### *RITMO1* affects fluorescence rhythmicity under free-running conditions

In order to characterise the role of RITMO1 within the diatom’s time-keeper machinery, we performed gene editing by proteolistic bombardment of the CRISPR-Cas9 protein (Serif *et al*., 2018). Sequence analyses confirmed the mutagenesis of *RITMO1* by biallelic frameshift mutations in two lines, defined as KO1 and KO2 (Fig. S4). The *P. tricornutum* wild-type (WT), the two KO lines, a transgenic line not presenting any mutation for *RITMO1* (CTR) and the previously generated *RITMO1* over-expressor OE1 line (Annunziata *et al*., 2019) were analysed regarding their fluorescence rhythms.

We first tested the effect of *RITMO1* KOs and overexpression in cells exposed to L:D cycles of moderate intensity (25 µmol photons m^−2^s^−1^) under various photoperiods (Fig. 2). Strong fluorescence oscillations were observed in all the lines and conditions tested (Fig. 2, Fig. S5 and Table S3), for cultures exhibiting between 0.8 division/day (under 8L:16D, and 12L:12D, Table S3) and 1 division/day (under 16L:8D). *RITMO1* KO and OE lines showed rhythmic fluorescence trends comparable to those of the WT and CTR lines (Fig. 2, Fig. S5 and Table S3).

**Fig. 2:**
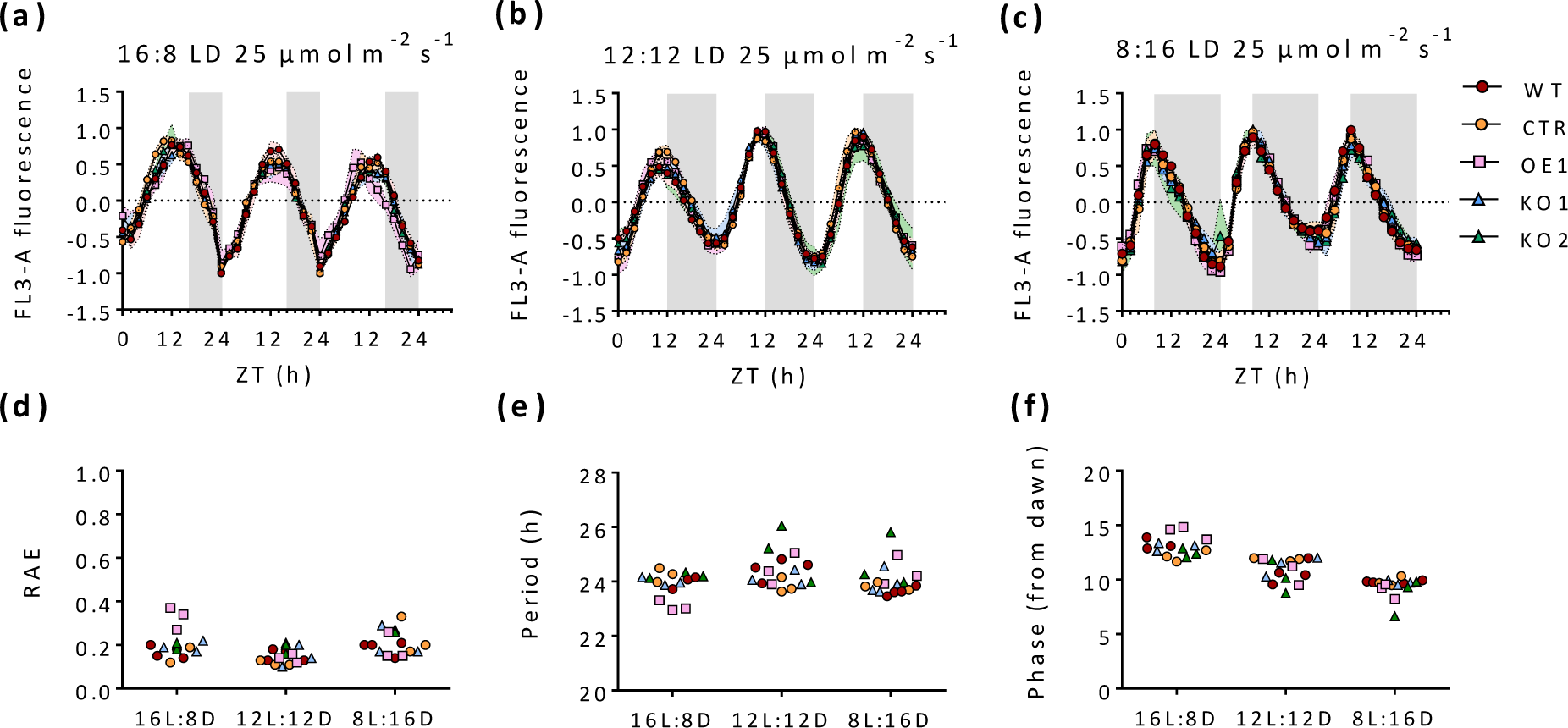
Characterization of cellular fluorescence rhythmicity of *P. tricornutum* WT, CTR, and *RITMO1* OE1, KO1 and KO2 lines (n>=3) under L:D cycles. Cellular fluorescence profiles (FL3-A parameter) of cells adapted to 25 µmol photons m^−2^ s^−1^ under different photoperiods: 16L:8D (a); 12L:12D (b); 8L:16D (c). Relative Amplitude of Error of FFT-NLLS fit (d); Period (e) and Phase estimation (relative to dawn) (f) calculated with FFT-NLLS algorithm. Fluorescence profiles were baseline detrended and then normalized between 1 and −1 for graphic representation. Colored dashed lines represent standard deviations. White and grey regions represent light and dark periods.

We then examined circadian rhythms in WT, CTR, *RITMO1* KO and OE lines following a shift from L:D to L:L (Fig. 3 and Fig. S5), applying the previously established conditions for detection of the circadian clock (25 L:D to 17 L:L µmol photons m^−2^ s^−1^). Strongly altered FL3 rhythmicity was observed in OE1 (Fig. 3 and Table S4) in L:L, confirming results obtained previously under constant blue light (Annunziata *et al*., 2019). Here, we showed that rhythmicity is also significantly affected in the KO lines (Fig. 3 and Table S4). In cases where we were able to detect rhythms in KO1 and KO2 in L:L, however, they showed a reduced amplitude, when compared to WT and CTR, which is paralleled by an RAE increase (Fig. 3b-e and Table S4). In these experiments, we also monitored the growth of the different cell lines (Fig. 3e, Fig. S6 and Table S4). All the strains showed a comparable growth, excluding that the alterations of circadian FL3 rhythmicity observed in RITMO1 KO and OE1 are attributable to growth defects. In addition, both KO lines and the OE line showed a similar alteration of rhythmicity under higher L:L light intensity (25 L:D to 25 L:L µmol photons m^−2^ s^−1^), while clock activity was still detectable in the WT and CTR strains (Fig. S7).

**Fig. 3:**
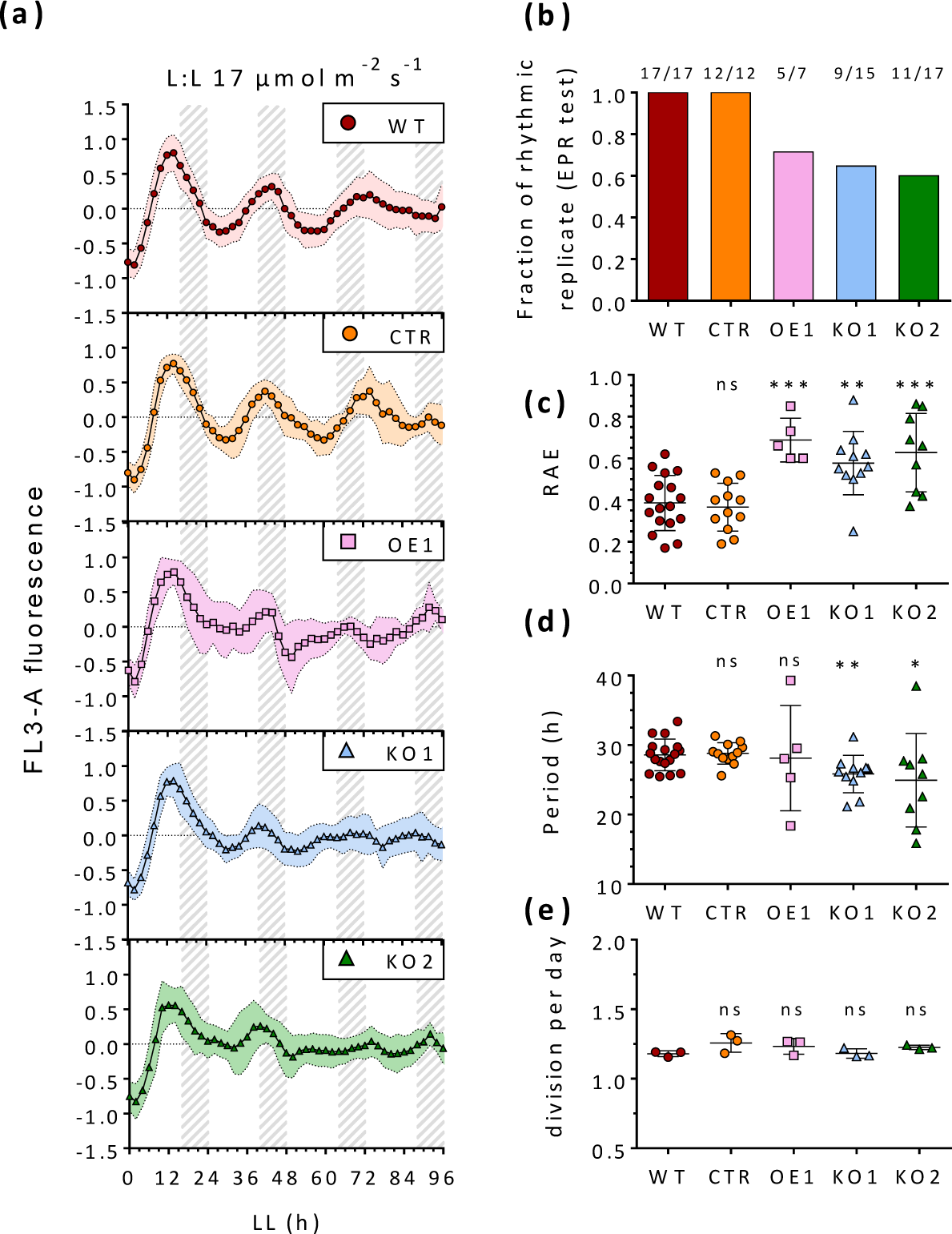
Analysis of the effect of *RITMO1* knock-out (KO) and ectopic overexpression (OE) on circadian fluorescence rhythmicity. (a) Fluorescence profile (FL3-A parameter) of cells adapted to 16L:8D 25 µmol photons m^−2^ s^−1^ and subsequently exposed to L:L 17 µmol photons m^−2^ s^−1^ in WT, CTR, OE1, KO1 and KO2 strains (n>=7). (b) Fraction of replicates that passes EPR test for all strains. Further analyses are only performed for the lines found rhythmic with EPR test: (c) Relative amplitude of Error of the FFT-NLLS method fit; (d) Period estimation obtained with FFT-NLLS method. (e) Division per day along the experiment (n=3). Fluorescence profiles were baseline detrended and then normalized between 1 and −1 for graphic representation. Colored dashed lines represent standard deviations. In L:L, the time is counted starting from the end of the last L:D day. Statistical differences were examined using unpaired Student’s t-test with the WT as reference sample (* = p<0.05; ** = p<0.01;*** = p<0.001).

We further inquired whether alteration of *RITMO1* expression influences the capacity of the cells to dynamically respond to environmental inputs. To do so, non-synchronized cells grown in continuous light for two weeks were exposed to a 16L:8D photoperiod (Fig. 4, Fig. S5d), in a mirror treatment to that reported in Fig 3. All the lines recovered 24 h fluorescence rhythmicity already after the first night and no phase shift was observed, indicating that a treatment of 8 h of darkness is sufficient to re-synchronize the population and that RITMO1 is not essential for photoperiodic entrainment of the FL3 rhythms.

**Fig. 4:**
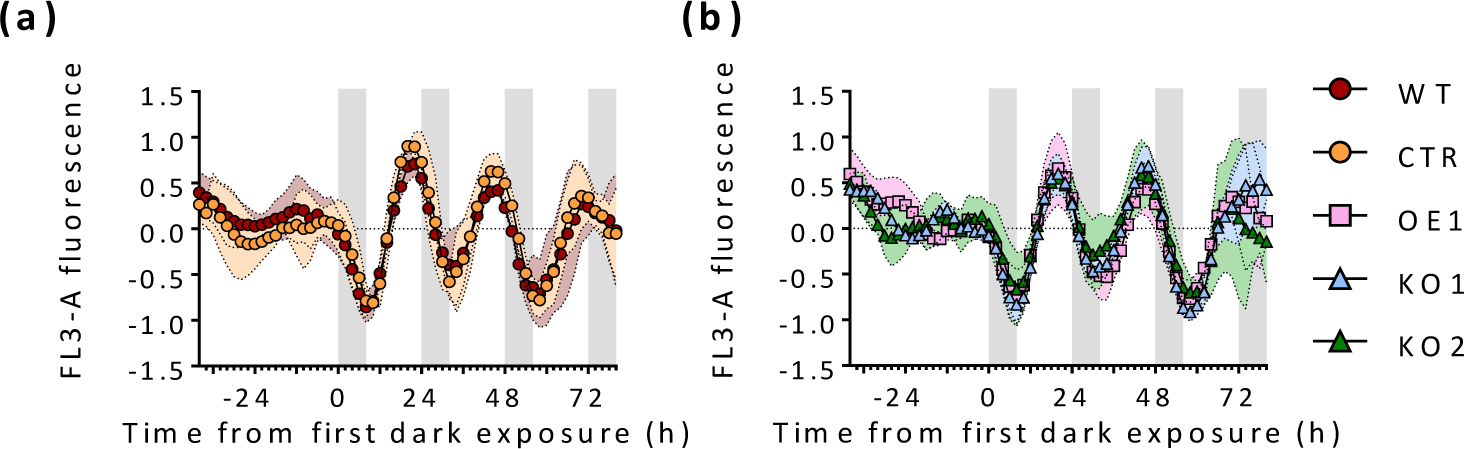
Characterization of cellular fluorescence (FL3-A parameter) rhythmicity of WT, CTR (a) and *RITMO1* OE1, KO1 and KO2 (b) strains (n>=3) re-exposed to 16L:8D cycles following constant light desynchronization. Fluorescence profiles were baseline detrended and then normalized between 1 and −1 for graphic representation. Colored dashed lines represent standard deviations. White and grey regions represent light and dark periods.

### RITMO1 is implicated in the circadian regulation of photosynthesis

The observed rhythmic patterns of FL3 in free running conditions could reflect rhythmic patterns in plastid content or/and activity, e.g., the number of PSIIs per cell or in the number of chlorophyll molecules per PSII (i.e., related to the functional absorption cross-section of PSII, σPSII). This prompted us to monitor several photosynthetic parameters with PSII fluorescence (Materials and Methods and Method S1) and to investigate the involvement of RITMO1 in circadian control of diatom photophysiology. The measured parameters comprise four groups of processes: i) PSII integrity and light absorption capacity (F_v_/F_m_ and σPSII), ii) photosynthetic electron transport, iii) maximal photoprotection capacity (NPQ_m_) and, iv) kinetics and regulation of NPQ (E50NPQ and NPQrel.), the latter in *P. tricornutum* being strictly mediated by the xanthophyll cycle (XC) (Blommaert *et al*., 2021). This analysis was first carried out on 16L:8D cells, where all measured parameters showed clear diel oscillations (see Fig. 5, Fig. S8 and Table S5). Cells were then transferred to L:L, and the photophysiology analysis was resumed on the second and the third days (avoiding transfer-related transient responses in the first 24 h) (Fig. 5a). Only few of the photosynthetic parameters maintained strong and statistically significant oscillation in free running L:L conditions in the WT (*p.value*<0.01 (Table S5)). The two observables informing on the dynamic regulation of the XC, through the light-dependence (E50NPQ) and kinetics in darkness (NPQrel.) of NPQ, showed the most pronounced rhythms, whereas a significant rhythmic behaviour for F_v_/F_m_ (was also observed (Fig. 5, Table S5). In the case of RITMO1 OE and KOs, values of these three parameters showed high dispersion between replicates and no rhythms were detected by statistical tests in L:L. In the case of OE lines, a slight phase anticipation effect was observed for NPQ dynamics parameters in L:D. These results support a role of RITMO1 in the circadian regulation of diatom photophysiology.

**Fig. 5:**
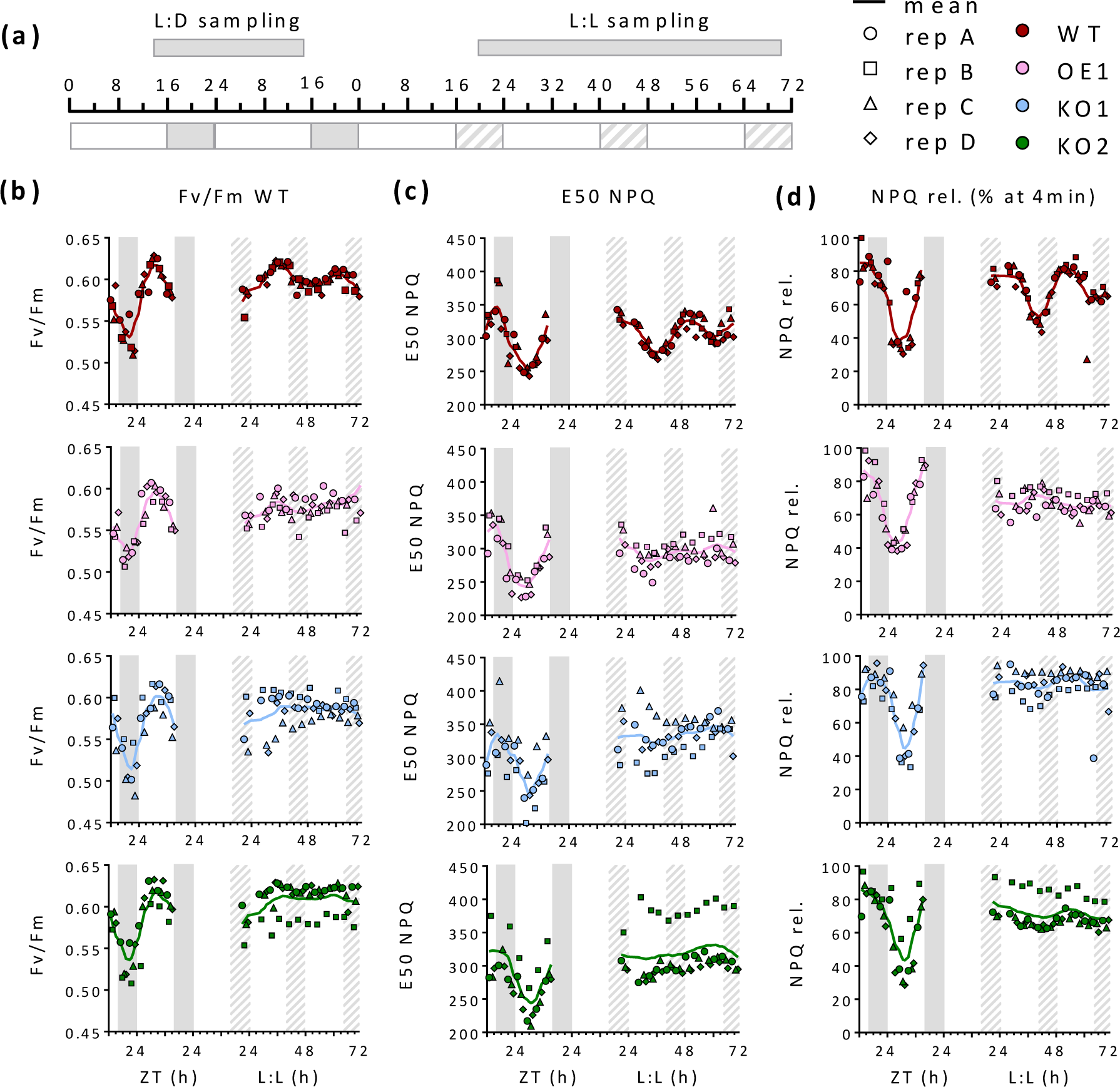
Analysis of the effect of *RITMO1* knock-out (KO) and ectopic overexpression (OE) on different photosynthetic parameters. (a) Schematic representation of sampling times for the analysis of WT, KO1, KO2 and OE1 cells grown under 16L:8D 50 µmol photons m^−2^ s^−1^ and subsequently exposed to L:L 30 µmol photons m^−2^ s^−1^. Samples were collected continuously and measured one by one in the following order WT, KO1, KO2, OE1 from replicate A to replicate D, with 15 minutes interval due to the measurement. Measurements were performed over 24 h in L:D and 48 h in L:L. In free-running, samples were collected starting at L:L 16. (b) Maximum dark-acclimated quantum yield of PSII (Fv/Fm), (c) E50NPQ, e.g., light intensity where 50% of the maximum NPQ is reached and (d) NPQ relaxation, *e.g*., percentage of NPQ at maximum light intensity relaxed after 4 minutes in the dark. Dots, squares, triangles and diamonds represents the individual measures for biological replicates A to D, lines represent the moving average of all replicates (window of 4 h). White and gray regions represent light and dark periods, gray dashed regions represent subjective nights in free-running conditions

### RITMO1 plays a central role in the circadian control of gene expression in diatoms

We investigated the role of RITMO1 in the circadian regulation of gene expression, by analysing the expression of a set of genes by RT-qPCR in cells entrained under 16L:8D and then transferred to L:L. Gene expression was measured every 4 h for two days in L:D. Under L:L, sampling was performed every 4 h between L:L32 to L:L68, corresponding to the second and third subjective night and day, as in the photosynthesis experiment (Fig. 6a). WT, OE1 and KO2 lines were chosen for the analysis. *RITMO1* showed rhythmic expression patterns in L:D, which are maintained in L:L in the WT, although with a delayed peak of expression (Fig. 6b). As previously reported (Annunziata *et al*., 2019), the OE1 line showed higher *RITMO1* expression levels and anticipation of the maximum of expression compared to the WT in L:D (Fig. 6b), due to the overexpression of *RITMO1* driven by the *FcpB* promoter. In L:L, OE1 displayed higher expression level than WT and a distinct rhythmic pattern, with an acrophase during the subjective light period. For KO2, an expression pattern similar to the WT was observed for *RITMO1* in L:D, while loss of circadian expression was observed in L:L. This indicated that the CRISPR-Cas mutagenesis did not significantly alter *RITMO1* mRNA expression in L:D, but generated a non-functional protein (Fig. S4), likely perturbating a circadian autoregulatory feedback loop. We also analysed other TFs transcripts (*bHLH1b*, *bHLH3* and *bZIP7*) exhibiting strong abundance rhythmicity in L:D cycles (Annunziata *et al*., 2019). Expression peak anticipation was observed for *bHLH3* in L:D in the OE1 only (Fig. 6c). Rhythms were maintained in L:L in WT cells (Fig. 6c), but they were strongly altered in both OE1 and KO2 for the *bHLH1b* and *bZIP7* transcripts. For *bHLH3*, a damped rhythmicity was still detected in KO2, although with an altered phase.

**Fig. 6:**
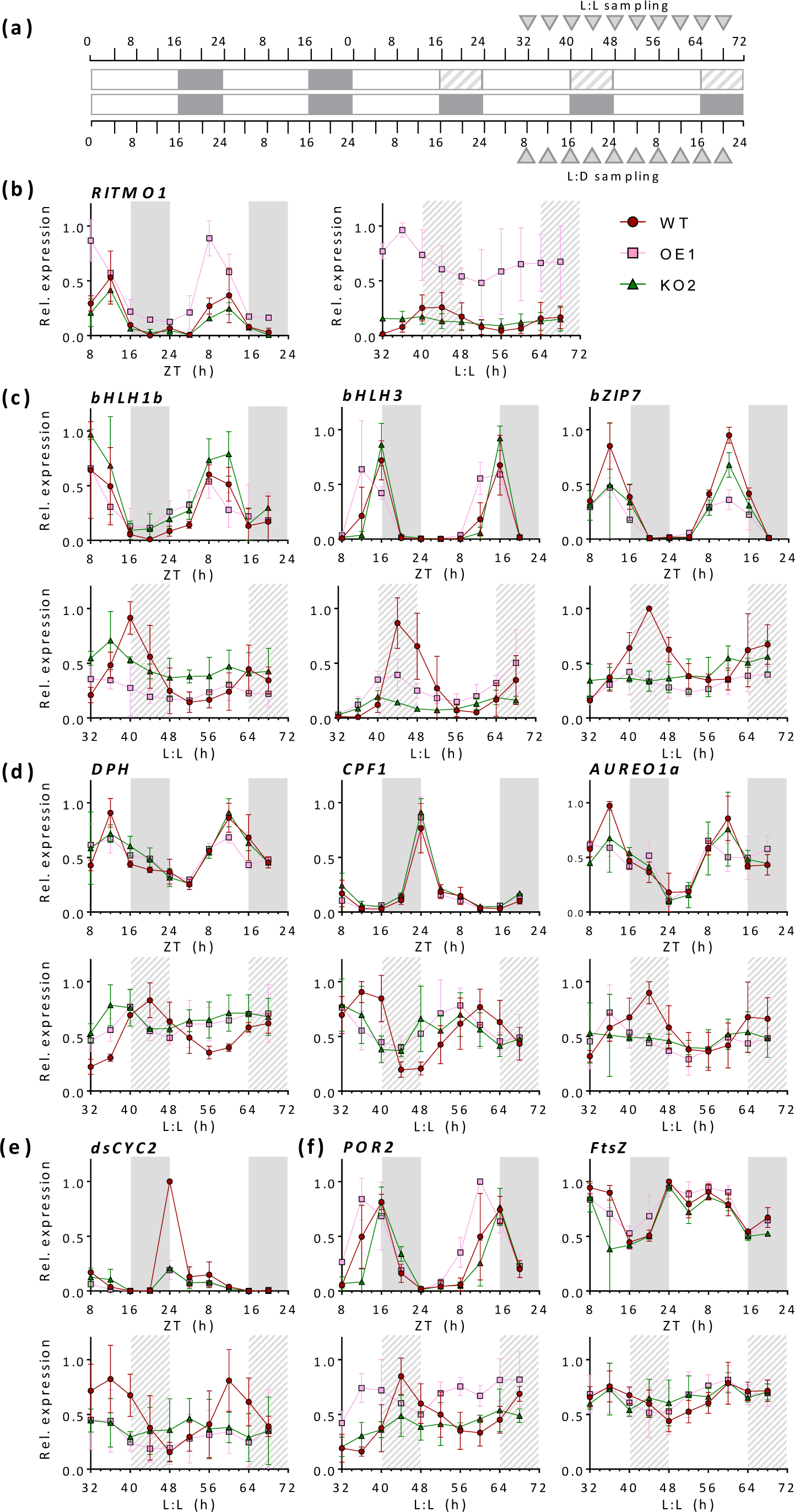
Analysis of the effect of *RITMO1* knock-out (KO) and ectopic overexpression (OE) on gene expression rhythmicity. (a) Schematic representation of the sampling points: samples were collected every 4 h over 36 h in parallel from cells adapted to 16L:8D cycles 25 µmol photons m^−2^ s^−1^ and from L:L 17 µmol photons m^−2^ s^−1^. In free running, samples were collected starting at L:L 32 (mean ± SEM,n = 3). Analyses by qRT-PCR of selected genes in WT, KO2 and OE1 lines: (b) *RITMO1* expression profiles of endogenous and ectopic *RITMO1* transcripts; (c) Transcription factors *bHLH1b*, *bHLH3* and *bZIP7* under L:D cycles (top) and L:L (bottom); (d) Photoreceptors diatom phytochrome (*DPH*), Cryptochrome Photolyase Family 1 (*CPF1*) and Aureochrome 1a (*AUREO1a*); (e) Expression profiles of the cyclin *dsCYC2*, (f) protochlorophyllide oxidoreductases 2 (*POR2*) and (g) filamenting temperature-sensitive (*FtsZ*) genes. Expression values represent the average of three technical replicates for each biological replicates (n=2 in L:D and n=3 in L:L) ±SD, normalized using the *RPS* and *TBP* reference genes. Expression values are given relative to the maximum expression for each gene, where ‘1’ represents the highest expression value of the time series. White and grey regions represent light and dark periods, grey dashed regions represent subjective nights in free-run conditions.

We next analysed the expression of diatom photoreceptors, potential actors in the light input pathway to the clock (Fig. 6d). Rhythmic expression was observed for the phytochrome photoreceptor (*DPH*) (Fortunato *et al*., 2016; Duchene *et al*., in press) in L:D and at the second and third day in L:L in WT. Only minor residual oscillations were observed in OE1 and KO2 at the second subjective day, compared to the WT. Rhythmic patterns in L:D and at the second and third L:L period were also detected for the blue-light receptor *CPF1*. KO2 and OE1 showed no alteration in L:D compared to WT, but a possible temporal shift when compared with WT in L:L. The pattern of the *AUREO1a* transcript was particularly interesting since this blue light receptor also acts as a TF controlling the expression of an important number of genes in *P. tricornutum*, including *RITMO1* under L:D and D:D conditions (Madhuri et al., 2024). The *AUREO1a* rhythmic expression pattern observed under L:D and L:L conditions in the WT was altered in *RITMO1* KO2 and OE1 in L:L, suggesting that that RITMO1 and AUREO1a might be part of a same regulatory system. This hypothesis is further supported by the altered circadian FL3 fluorescence rhythmicity observed in *AUREO1a* KO mutants, compared to WT cells in L:L (Fig S9).

We then investigated the expression of putative clock regulated genes. The cell cycle regulator *dsCYC2*, whose expression is strongly and transiently induced by the blue light-receptor AUREO1a after switching from darkness to light (Huysman *et al*., 2010, 2013) showed a rhythmic expression in the WT in L:D, with an increase in the expression before the light onset (Fig. 6e), which is lost in OE1 and KO2. In LL, *dsCYC2* rhythmic expression pattern was observed for WT, but not for the OE1 and KO2 (Fig. 6e). The *CYCP6* gene, whose expression has been shown to be associated with the G1 phase, had a rhythmic expression in L:D, with a peak at the end of the night, in all the analysed strains. Minor rhythmicity was observed in WT in the second free running day, further reduced at the third day. Rhythmic patterns were not observed in the transgenic KO and OE lines for *CYCP6* (Fig. S10). The *CYCB1* gene, considered as a marker of the G2/M phase, showed a clear rhythmic pattern in L:D, peaking at the end of the light period, which was overall unaltered in KO2 and OE1. However, the expression of *CYCB1* appeared arrhythmic in L:L in all the lines (Fig. S10).

Finally, we analysed the expression patterns of two genes linked to chloroplast biogenesis and activity: *FtsZ*, encoding a putative component of the diatom chloroplast division machinery (Gillard *et al*., 2008; TerBush *et al*., 2013) and *POR2 (*protochlorophyllide oxidoreductase 2) putatively involved in chlorophyll biosynthesis (Hunsperger *et al*., 2016) (Fig. 6f). For *POR2*, we observed rhythms in both L:D and L:L. Gene expression anticipation was observed for the OE in L:D, while deregulation of rhythmic pattern was observed for both OE and KO in L:L. On the contrary, for *FtsZ* we observed rhythmic oscillation in L:D, but not in L:L, even for the WT. Interestingly the expression pattern is not altered in *RITMO1* OE or KO lines, ruling out the possible regulation of *FtsZ* by an endogenous oscillator.

### RITMO1 mutation complementation rescues circadian rhythms in *P. tricornutum*

To further prove the involvement of RITMO1 in circadian rhythms, we complemented the KO2 line with a construct containing the genomic *RITMO1* sequence fused to the *Venus* reporter under the control of *RITMO1* promoter and terminator sequences (Method S3). Different transgenic lines were obtained and screened by Western blot by using anti-GFP antibody to check transgene expression time and levels at the beginning and end of the light period (Fig. S11a). Nuclear localization was confirmed for these lines by fluorescence microscopy (Fig. S11b). However, when we analysed FL3 rhythms, only the KO2-C2 line, which display moderate and rhythmic expression of RITMO1 (S11a), rescued the WT circadian patterns (Fig. 7a). We further use this line to analyse gene expression rhythms. A specific feature of the circadian clock is the capacity to anticipate periodic L:D changes. In Fig 6 we observed lower expression of the cell cycle regulator *dsCYC2* in KO and OE, compared to WT, at the D to L transition. Therefore, we analysed *dsCYC2* expression with a higher temporal resolution before and after the light onset (Fig. 7b). The analysis indicated a delay in the light anticipation response and in the rapid-light induced expression in KO2 compared with WT (Fig. S7b) that was rescued in the KO2-C2 line. Analyses of selected genes in L:L also confirmed a recovery of rhythmic expression pattern in free run for all the analysed genes. For some genes, such as *AUREO1a* and *bZIP7* we observed an anticipation of the expression time in KO2-C2, and for *dsCYC2* (Fig. S12), a reduced amplitude. These phenotypes could be linked to a partial deregulation of the regulatory loop involving RITMO1 due to the gene over-expression in the KO2-C2 line compared to the WT (Fig. 7c) or deregulation of still uncharacterized RITMO1 interactors.

**Fig. 7:**
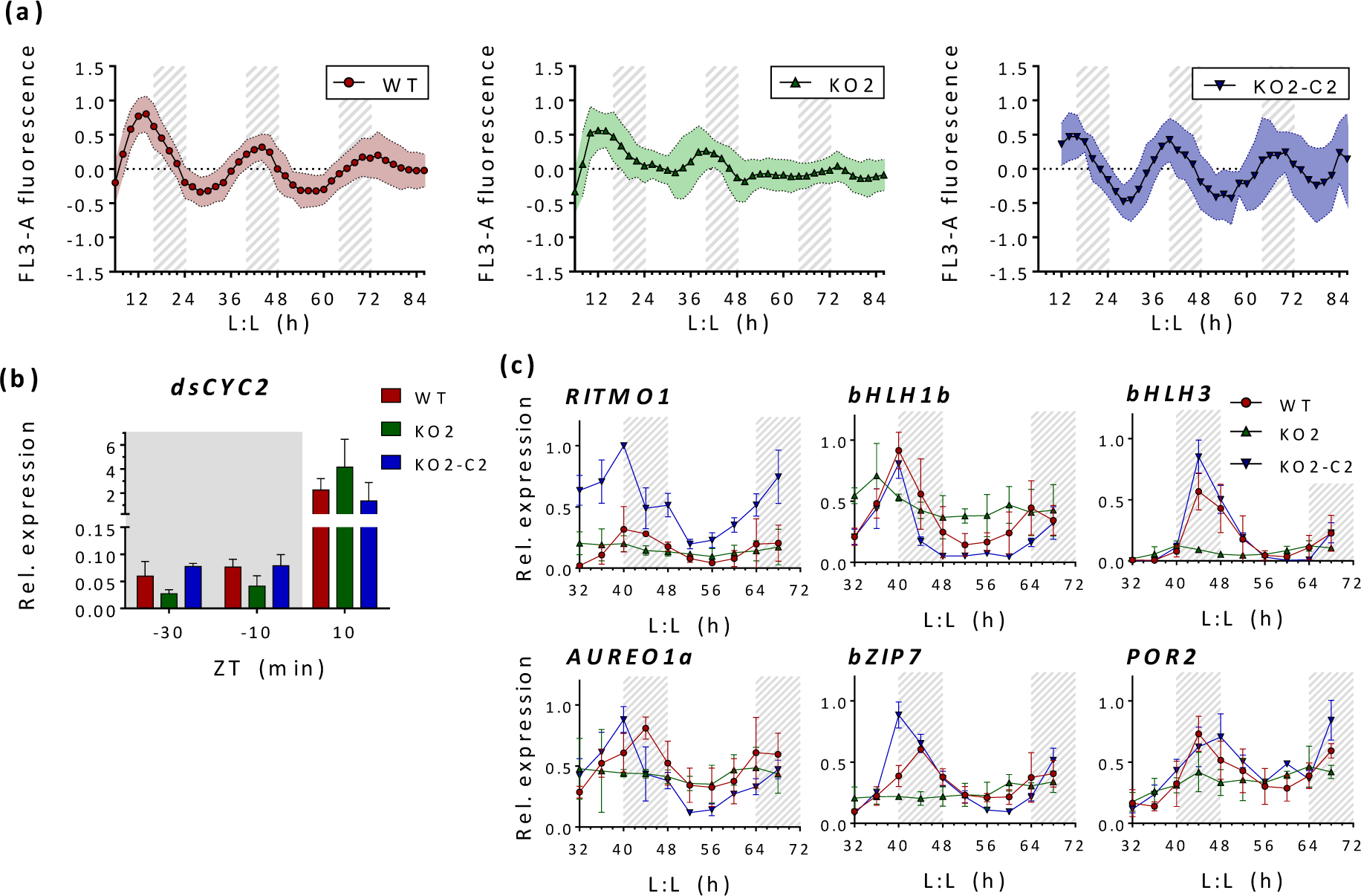
Analysis of rhythmicity in *RITMO1* knock-out strain (KO2) transformed with the RITMO1p:RITMO1VENUS:RITMO1t construct to obtain the KO2-C2 complemented line. (a) Normalised and baseline detrended cellular FL3-A fluorescence profiles in free run from WT, KO2 and complemented KO2-C2 strains (n>=6), entrained at 16L:8D 25 µmol photons m^−2^ s^−1^ before the shift to L:L 17 µmol photons m^−2^ s^−1^. (b) Analysis by qRT-PCR of *dsCYC2* relative expression in WT, *RITMO1* KO2 and KO2-C2 lines grown under 16L:8D cycles 25 µmol photons m^−2^ s^−1^ and collected for the analysis before and after the light onset at the time indicated. Red, green and dark blue bars represent mean values for WT, KO2 and KO2-C2 respectively (n=3) ±SD, normalized using the *RPS* and *TBP* reference genes. (c) Analyses by qRT-PCR in L:L of the expression profiles of selected genes in WT, KO2 and KO2-C2 lines: endogenous and ectopic *RITMO1* transcripts; *bHLH1b*, *bHLH3* and *bZIP7*, *AUREO1a* and *POR2* (three biological replicates ±SD, normalized with using the RPS and TBP reference genes. Expression values are given relative to the maximum expression for each gene, where ‘1’ represents the highest expression value of the time series. White and grey regions represent light and dark periods, grey dashed regions represent subjective nights in free-run conditions.

## Discussion

The characterization of various biological rhythms in this study reveals a complex regulation of diatom responses to light and dark cycles, involving both environmental response pathways and internal biological programs. This was established by identifying experimental conditions to monitor the circadian clock activity and the effects of its perturbation on various rhythmic processes In the case of cell fluorescence, where a large quantity of data was obtained over several days and under different conditions, we observed the persistence of rhythmicity for at least 4 days, in the absence of exogenous temporal cues. This confirms the existence of an endogenous timekeeper in diatoms. In free running, the periods of FL3 are overall longer than 24 h (between 26 and 28 h) at the different light conditions tested, indicating that the circadian clock free-runs with its own period, which does not match exactly 24 h, a typical feature of circadian rhythms (McClung, 2006). As in other organisms, the diatom circadian rhythms exhibit temperature compensation below the growth temperature and up to 2°C above (Fig. 1 and Fig. S3). The amplitude of circadian oscillations also depend on the light intensities experienced by the cells during the entrainment and free-running conditions (Oakenfull & Davis, 2017; Webb *et al*., 2019). The most robust circadian rhythms were observed for cells exposed to moderate light intensity, resulting in an average of one division per day. On the contrary, limited persistence of circadian rhythmicity was observed under very low light, when light intensities are high enough to promote a more sustained growth (Fig 1) or at temperatures higher than 20°C, likely affecting the metabolism (Fawley, 1984; Rehder *et al*., 2023). Poor free-running rhythmicity could result from a loss of rhythmicity or from asynchrony among individual cells within the bulk culture. *De facto*, FL3 is a good proxy of the synchronicity in diatom population as it captures different observables, including cell growth, cell cycle progression (Huysman *et al*., 2010) and plastid biogenesis and activity (Falciatore *et al*., 2022). However, all these processes are also depending on direct environmental inputs, which could override clock activity. Moreover, darkness provokes cell cycle arrest in diatoms, which explains the loss of FL3 rhythmicity in D:D. The forced desynchronization experiments using 6L:6D cycles, normally used to elucidate circadian clock activity, leads to short oscillations occurring twice per circadian cycle in *P. tricornutum* and also affect diatom growth (Fig. 1). Our FL3 analyses suggests a temporal control of the diatom cell division, which appears to be limited to a window of time, as shown in many organisms, including the green algae Chlamydomonas (Goto & Johnson, 1995), *O. tauri* (Moulager *et al*., 2007, 2010) or *Nannochloropsis* (Braun *et al*., 2014; Poliner *et al*., 2019). In these algae, cell division occurs when the cells have reached a critical size defined by the metabolic status and are in the time window defined by the circadian clock (timer). The gating of the cell cycle by an endogenous oscillator has not been demonstrated in our study, but it is possible to hypothesize that the circadian clock and the cell cycle co-act to control light-dependent regulation of diatom cell division, as suggested by the regulation of the *P. tricorntum* cell cycle regulator *dsCYC2* by RITMO1 (Fig. 6, Fig. S7 and below) and AUREO1a (Huysman *et al*., 2013). The perturbation of circadian rhythmicity of *P. tricornutum* fluorescence may therefore arise from uncoupling of circadian control of cell division from direct response to changing environmental cues, as shown in *O. tauri* under strong light (Moulager et al., 2007). Additionally, acclimation of photosynthesis could also play a role. In many phototrophs, photosynthesis is a major circadian output, and our data also support its circadian regulation in diatoms (see below). Evidences indicate that photosynthesis can, in turn, regulate clock activity by acting on input pathways (Farré & Weise, 2012; Haydon et al., 2017; de Barros Dantas et al., 2023). Therefore, it is likely that, in addition to the cell cycle, a strong interplay exists between the circadian clock and photophysiology, coordinating critical diatom cellular and metabolic activities under variable environmental conditions. These aspects will deserve further characterization via the generation of ad hoc mutants in distinct light-regulated processes and extended physiological characterization, as has been elegantly shown in plants (Haydon et al., 2013).

Identifying the specific conditions under which clock activity can be reliably observed has been essential for extending the analysis of circadian regulation to relevant physiological processes and further investigating the diatom circadian clock system through RITMO1. Here we show that all the circadian processes described in this study are affected not only by the ectopic over-expression of *RITMO1* but also by its mutagenesis (Fig. 3, Fig. 5 and Fig. 6, Fig. 7). The strong perturbation of various circadian rhythms by the knock-out of the single *RITMO1* gene, along with the rescue of circadian phenotypes through genetic complementation, unambiguously supports a role of this protein in the diatom circadian clock. Given the still limited information on molecular clock(s) in diatoms and other Stramenopiles, it is not yet possible to confirm whether RITMO1 is a component of the central oscillator or a slave oscillator (e.g., Heintzen *et al*., 1997), transducing the master clock’s impulses to a subset of circadian-regulated processes. Nevertheless, our data support that this factor with bHLH-PAS domains is part of a circadian regulatory loop generating rhythms: RITMO1 affects its own expression, that of other clock components such as AUREO1a and of output genes. As observed by the characterization of the *dsCYC2* gene expression, it also controls light anticipation responses, attenuated in the *RITMO1* mutant. Alteration of rhythmicity in L:D cycles is also observed for several genes in OE lines. The finding that circadian rhythms in KO lines are rescued only in the line showing the correct expression time and moderate gene over-expression compared to the WT, strongly support that the missregulation of RITMO1 might have a stronger effect than gene obliteration on the transcriptional feedback loops generating rhythms, as reported for OtCCA1 (Corellou *et al*., 2009).

Other TF members of the bHLH family (*bHLH1b* and *bHLH3*), as well as *bZIP7*, represent direct or indirect RITMO1-target genes possibly involved in the circadian clock (Fig. S14). A molecular interplay in circadian regulation can now be proposed for AUREO1a and RITMO1, two TFs showing similar expression time at the end of the light period. *RITMO1* expression is regulated by *AUREO1a* (Mann *et al*., 2020; Madhuri *et al*., 2024) and we here showed that the circadian expression of *AUREO1a* is altered in *RITMO1* mutants, and that both factors are essential in sustaining circadian fluorescence rhythmicity. The rapid re-entrainment of biological rhythms in both desynchronized WT and *RITMO1* mutants when re-exposed to the light dark cycles suggests that RITMO1 is not essential for clock entrainment (Fig. 4). However, this function could be expected of AUREO1a because of its additional function as blue light photoreceptor. The circadian regulated photoreceptors Phytochrome DPH and the blue light Cryptochrome Photolyase 1 CPF1 are additional candidates of the diatom input pathways (Fig. 6). Further investigations under different fluence rate/ quality of light and phase response curves in the confirmed and putative diatom clock components (Fig. S14) would help to further assess the role of these factors in the diatom circadian clock system and their possible interaction with the photic control of circadian rhythms.

In addition to gene expression, here we demonstrate that circadian clock controls several photosynthetic parameters in *P. tricornutum*, as observed in plants (Dodd *et al*., 2005; Häfker *et al*., 2023) and other algae (Sorek *et al*., 2013) and supporting previous observations based on the analysis of carbon uptake or pigment content in diatoms (Palmer *et al*., 1964; Ragni & d’Alcalà, 2007). Parameters informing on photosynthetic electron transport (maximal capacity under saturating light or performance under light-limiting regime) or photoprotection capacity (NPQ_m_) show diel cycles but do not significantly oscillate under free running conditions (Fig S6, Table S5). Although we cannot rule out that these parameters display oscillatory patterns below the sensitivity of our methodological approach, these rhythms are clearly smaller than the ones linked to PSII light absorption and conversion (Fv/Fm and σPSII) and especially to the regulation of NPQ (E50NPQ and NPQrel.), which are under RITMO1-dependent circadian control. Many known rhythmic processes can influence Fv/Fm, such as PSII recycling (Li *et al*., 2016) or NPQ relaxation (this study). The circadian regulation of chlorophyll biosynthesis (Ragni & d’Alcalà, 2007; Hunsperger *et al*., 2016), here supported by the regulation of the *POR2* gene by RITMO1 (Fig. 6 and 7), could influence circadian rhythms of σPSII.

Interestingly, the most significantly photosynthetic parameters under circadian control are the ones informing on the dynamics of fast NPQ regulation. The ecological success of diatoms in turbulent waters is often attributed to their efficient photoprotective capacity via NPQ (Lavaud *et al*., 2007), which allows avoidance of photodamages under high light, while being rapidly tuned down to avoid “energy wasting” when irradiance decreases. Therefore, it is tempting to suggest that clock regulation of NPQ dynamics contributes in the real-time adjustment/anticipation of photoprotective mechanisms to the light environment and time of the day. The absence of persistence of oscillations in the maximal NPQ capacity (NPQ_m_) in free-running makes it unlikely that the concentration of the main NPQ molecular actors, the Lhcx proteins and the xanthophyll cycle (XC) pigments (Buck *et al*., 2021, Croteau *et al*., 2024), are a primary target of the circadian clock. At the contrary, the circadian regulated E50NPQ and NPQrel. parameters only depend on the relative rates of the diadinoxanthin de-epoxidase (increasing NPQ) and diatoxanthin epoxidase (decreasing NPQ) enzymes of the XC (Blommaert *et al*., 2021). As some XC genes maintain rhythmic expression in the dark (Annunziata *et al*., 2019), RITMO1 could contribute to circadian regulation or to post-translational modification of these enzymes. Alternatively, RITMO1 could regulate the availability of co-substrates of the XC enzymes, luminal ascorbate for diadinoxanthin de-epoxidase or stromal NADPH for diatoxanthin epoxidase, which in turn, could impose circadian rhythms in the kinetic rates of those enzymes, regardless of their quantity. Circadian rhythms of NADPH levels are known in *A. thaliana*, as well as in ascorbate pathway genes in plants (Dowdle *et al*., 2007) and *Euglena gracilis* (Kiyota *et al*., 2006). Interestingly, the anticipation for NPQrel. observed in OE1 line in L:D compared to WT and KO lines (Fig. 5d), suggest that even under enforced photoperiodic rhythms, RITMO1 accumulation may modulate the activities of the XC enzymes and NPQ dynamics.

Finally, this study establishes marine diatoms as powerful experimental models for research on chronobiology in secondary endosymbionts, which form the bulk of phytoplankton. Future research in these organisms will help addressing important questions regarding the evolution and functional diversification of circadian systems (Laosuntisuk *et al*., 2023) and to assess the importance of the circadian clock for life in the Oceans. The *P. tricornutum* RITMO1 mutants show no growth defect in either L:D or in L:L. At this stage, we do not know if this is due to the potentially redundant function of other clock proteins (e.g., other bHLH-PAS) or to the peripheral role of RITMO1 in the circadian system, or to the weak activity of the diatom clock (Bordyugov *et al*., 2015; Poliner *et al*., 2019), which may not be necessary for optimal growth under laboratory conditions. In cyanobacteria, the effect of clock mutation on fitness is only clearly seen in competition experiments between circadian period mutants and wild-type cells and when their internal rhythms match those of the environmental cycle (Ouyang *et al*., 1998; Woelfle *et al*., 2004). Similar observations are also made with a variety of clock mutants in plants (Dodd *et al*., 2005; Yerushalmi *et al*., 2011). Therefore, further research with multiple clock mutants exposed to different environmental L:D cycles will be needed to clearly assess the role of the diatom clock on fitness.

The perturbation of circadian rhythms observed in *P. tricornutum* already at 2°C above the temperature of growth, similar to what reported for the green alga *Nannochloropsis oceanica* (Poliner *et al*., 2019) raises questions on the impact of temperature on the clock of phytoplankton. These organisms in the ocean may experience much less drastic temperature changes during L/D cycles compared to terrestrial organisms or marine organisms leaving in intertidal zones. Therefore, a more detailed study of the effects of temperature variations on circadian processes, as well as on the ability of temperature cycles to entrain the clock, will provide valuable insights into the significance of so far generalized clock properties for diatom biology. This understanding is crucial for accurately evaluating the impact of temperature changes on phytoplankton dynamics (Häfker *et al*., 2023). Notably, an expansion of bHLH-PAS TFs with different rhythmic expression profiles has been described in the benthic diatom *Seminavis robusta*, likely reflecting an adaptation and diversification of this protein family to both diel and tidal cycles (Bilcke *et al*., 2021). Thus, studies on rhythms and clocks in benthic diatoms could enhance our understanding of phytoplankton physiology in response to the various timescale cycles typical of the marine environment (Häfker *et al*., 2023). Further analysis of diatom responses to photoperiodic conditions is also warranted, as they dominate in polar regions, which are characterized by dramatic seasonal changes in photoperiod (Pierella Karlusich *et al*., 2020). The circadian clock play a central role in photoperiodic regulation in terrestrial plants and animals (Nishiwaki-Ohkawa & Yoshimura, 2016).

## Acknowledgements

This work has been supported by funding from the Fondation Bettencourt-Schueller (Coups d’élan pour la recherche francaise-2018), the “Initiative d’Excellence” program (Grant “DYNAMO,” ANR-11-LABX-0011-01), the ANR-DFG Grant “DiaRhythm (KR1661/20-1) to A.F. and P.G.K, and ANR Grant ClimaClock (ANR-20-CE20-0024) to A.F and F-Y.B, by the CNRS through the MITI interdisciplinary programs to M.J. D.C and B.B acknowledges the support by the European Research Council (ERC) PhotoPHYTOMIX project (grant agreement No. 715579). We thank Stephane Eberhard for critical suggestions and discussion.

## Author contributions

AF coordinated the project and designed research with AM and RM, and with the contribution of JPB, BB, MJ, FYB, and PGK; AM, RM, SCN, DC, LC, AH, NFS, BB, DC, DJ performed the research; AF, AM, RM, BB, JPB, FD, JYB, MJ, DC analysed and interpreted the data, AF, AM, RM wrote the manuscript. All the authors discussed the results and commented on the manuscript.

## Data availability

The original contributions presented in the study are included in the article and in the Supporting Information. Further inquiries can be directed to the corresponding author.

## Supporting Information

### Method S1

#### Analysis of photosynthetic parameters

Analysis of different photosynthetic parameters were performed by using a thorough PSII fluorescence protocol on cells grown as discussed in Materials and Methods.

Fluorescence induction from a saturating single turnover flash (100 µs) of blue light (450 nm) applied on dark-acclimated samples was used to calculate the functional absorption cross-section σPSII and connectivity of PSII (Gorbunov *et al*., 2020).

The light adapted φPSII was calculated as (Fm’ - Fs) / Fm’, where Fm’ represents the maximum fluorescence in light-adapted conditions and Fs is the steady-state fluorescence in the light. The relative electron transport rate (rETR) was then calculated as rETR = φPSII × E. The relationship between rETR and the light intensity (E) was fitted using the equation rETR = rETRm × (1 – exp (-α × E / rETRm)), where rETRm is the maximum rETR and α is the light-limited slope of the rETR vs E curve. Only rETR values from 25 and 50 µmol photons m^−2^ s^−1^ were used for the fit due to noise in the values above 600 µmol photons m^−2^ s^−1^.

The photoprotection capacity through Nonphotochemical quenching (NPQ) was calculated as Fm/Fm’-1, where Fm’ is the light acclimated maximal fluorescence, and the light dependence of NPQ was fitted as NPQ = NPQm × (E /(E50NPQ + E)*n, where NPQm is the (asymptotic) maximal NPQ capacity, E50NPQ is the light irradiance at half saturation of NPQ, and n is the sigmoidicity coefficient (Serôdio & Lavaud, 2011). Kinetics of NPQ relaxation were also investigated, which strictly reflects the activity of the xanthophyll cycle in *P. tricornutum* (Blommaert *et al*., 2021). Due to time constraints imposed by the experimental design, we only measured the first 4 min of the NPQ relaxation after the transition from the highest light step (where NPQ value is NPQ800) to darkness. From there, “NPQrel.” was calculated as (NPQ800 - NPQ4min) / NPQ800, where NPQ4min is the remaining NPQ after 4 min of relaxation (Fig. S1).

### Method S2

#### Cell cycle analysis

Cell cycle analysis was performed in cells grown under 16L:8D cycles. Around 1-2 10^6^ cells in exponential growth were collected by centrifugation every 3h for 24h. Cell pellets were fixed in cold 70% EtOH and stored for at least 24h in the dark at 4°C until processing. Fixed cells were then washed once in cold EtOH 70%, once in PBS and stained in the dark for 45 min at room temperature with DAPI (4’,6-diamidino-2-phenylindole) at a final concentration of 0.5ng ml^−1^. After staining, cells were washed with PBS and 50,000 events per sample were analysed by flow cytometry using the MACSQuant flow cytometer. The V1-A channel (408 nm excitation, 450/50 nm detection) was used to evaluate the content of DAPI-stained DNA. 2c and 4c peaks were used to infer the fractions of cells in different cell cycle phases, using a dedicated R script (Agier & Fischer, 2016).

### Method S3

#### Complementation of RITMO1 knock-out strain

##### U loop assembly

The uLoop modular cloning approach was used to generate, according to Pollak et al. (2020), a vector containing the genomic *RITMO1* sequence fused to the Venus reporter (*RITMO1:Venus*) with its endogenous promotor and terminator, and a resistance cassette to nourseothricine with *FcpB* promotor and *FcpA* terminator. RITMO1 promotor (*RITMO1p*, - 824 nt upstream start codon) and RITMO1 open reading frame (*RITMO1g*,Phatr3_J44962) blocks were newly synthetized by IDT, and the RITMO1 terminator (*RITMO1t*, +539 nt downstream stop codon) block was obtained after two rounds of PCR reaction with the primer pairs TermbHLH1.F.Rv and TermbHLH1.E.Fw, and UNS1FLFw and UNSXFLRv (Table S1). Fragments were assembled by Gibson in *Sap*I-opened pL0 as pL0_AC-RITMO1p, pL0_CD-RITMO1g and pL0_EF-RITMO1t, and subsequently assembled into pL1 pCAo2 along with Venus tag pL0_DE-Venus (obtained from Pollak et al, 2020). Nourseothricine resistance (*NrsR*) cassette was generated as pL1 pCAo2-1 vector by assembling FcpB promotor pL0_AC-FcpB (*FcpBp*) (obtained from Pollak et al. (2020)), and nourseothricine resistance gene (*NrsR*) pL0_CE_NAT and FcpA terminator pL0_EF-FcpAt (*FcpAt*) (obtained after two rounds of PCR with the NAT.C.Fw and NAT.E.Rv, and the FcpAt.E.Fw and FcpAt.F.Rv primer pairs, respectively, and UNS1FLFw and UNSXFLRv primers, and Gibson cloning into SapI-opened pL0) into pL1 pCAo1. RITMO1 fusion expressing vector and resistance cassette were finally assembled with pCAo3 and pCAo4 spacers in pL2 pCAe1 to obtain the final sequence that was transformed by biolistic in KO2 background. Map of this pL2 is presented in Fig. S14.

##### Screening Process

After transformation by biolistic method (Falciatore et al., 1999), an initial screen was performed on selective plates containing 50% F2 medium supplemented with 300 µg/mL nourseothricine. A second screen was conducted on plates with 95% F2 medium, also containing 300 µg/mL nourseothricine. The presence of the transgene in the transformed cells was confirmed by PCR (primer sequences listed in Table S1). On selected positive colonies, the expression of the RITMO1:Venus fusion protein was characterized at ZT1 and ZT11 under 12 hours light:12 hours dark (12L:12D) conditions by Western blot (Fig. S11). These strains were further tested for cellular fluorescence rhythmicity. One strain, KO2-C2, successfully rescued the cellular fluorescence phenotype under continuous light (L:L) conditions and was chosen for further gene expression analyses under L:D and L:L conditions via quantitative PCR (qPCR).

##### Immunoblot analysis

Proteins were extracted as previously described in Coesel et al. (2009). Membranes were blocked with PBS-T 5% milk (w/v) and incubated overnight at 4° C with α-GFP tag Mouse McAb (Proteintech) (1:5000) antibody. Following incubation with HRP-conjugated 521 secondary antibodies (Promega), proteins were detected with ClarityMax reagents (Bio-Rad) and imaged with a ChemiDoc Touch imaging system (Biorad, US).

##### RITMO1:Venus subcellular localization

The KO2 line complemented with the RITMO1p:RITMO1VENUS:RITMO1t construct where grown under L:D cycles and collected 1 h before the Dark period for microscopy analysis. For chlorophyll autofluorescence and Venus fluorescence cells were excited at 510 nm and detected at 650–741 nm and 529–562 nm respectively. Hoechst 33342 (Life Technologies) was used at a final concentration of 5µg/ml to stain nuclear DNA and stained cells were visualized by illumination at 405 nm and detection at 424–462 nm.

**Table S1:**
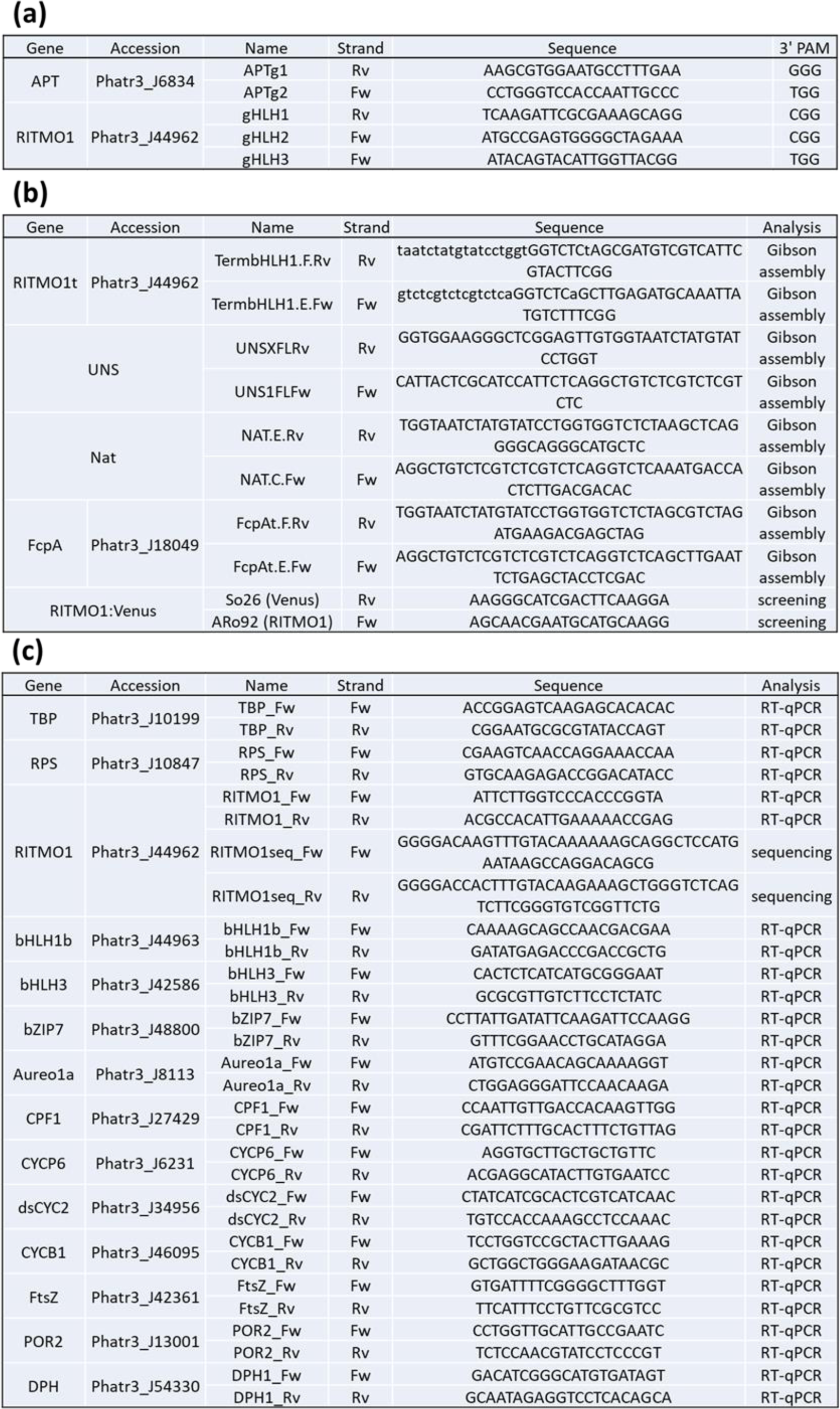
List of the spacer sequences for gRNAs and oligonucleotides used in this work. (a) List of the spacer sequences for gRNAs used for CRISPR-Cas9 mutagenesis of the *RITMO1* gene. Target gene, gRNA name, target strand, spacer sequence for gRNA and the associated protospacer adjacent motif (PAM) are reported. (b) List of the primers used for Gibson assembly of RITMO1t uLoop block and primers used for the screening of KO2 complemented strains (Method S3). (c) List of the oligonucleotides used for the sequencing of *RITMO1* WT and KO sequences and for RT-qPCR. Target gene, ID number in the Diatomicsbase (https://www.diatomicsbase.bio.ens.psl.eu/) (Villar *et al*., 2024), pairing strand and sequence of the oligonucleotides used in this work. Sequencing primers for *RITMO1* are allele specific.

**Table S2:** Rhythmicity parameters and growth of *P. tricornutum* cells grown under different L:D cycles of different light intensities and periods and following a shift to different free running conditions. Data in the upper table refer to WT cells entrained under 16L:8D 25 µmol photons m^−2^ s^−1^ at 18°C and subsequently exposed to different free-run conditions over four subjective days (D:D, L:L 5 µmol photons m^−2^ s^−1^, L:L 17 µmol photons m^−2^ s^−1^, L:L 25 µmol photons m^−2^ s^−1^) or to cells entrained under 16L:8D 75 µmol photons m^−2^ s^−1^ and after transition to L:L of 50 and 75 µmol photons m^−2^ s^−1^. Last column refers to data from cells entrained in 12L:12D 25 µmol photons m^−2^ s^−1^ and subsequently exposed to 6L:6D 25 µmol photons m^−2^ s^−1^. Data in the lower table refer to WT cells entrained under 16L:8D 25 µmol photons m^−2^ s^−1^ at 18°C and subsequently exposed to L:L 17 µmol photons m^−2^ s^−1^ at different temperature ranges: 14°C, 16°C with a temperature shift at the last light/dark transition and 20°C and 22°C with a temperature shift at the last dark/light transition. Rep. number indicates the number of biological replicas used for the analysis. EPR rhythmic lines indicates the lines that passed the EPR rhythmicity test for each condition and that are used for the FFT-NLLS rhythmicity test below. The FFT-NLLS algorithm was used to get the period, phase, amplitude, the Relative Amplitude of Error (RAE) that represents the amplitude of the error between the fit used by the algorithm and the data divided by the amplitude of the oscillations. The cell division/day was calculated in cells in exponential phase over the course of four days (n=3) for all lines (including the lines found non rhythmic by EPR analysis). Statistical differences were examined using unpaired Student’s t-test with the WT as reference sample.

| | 16L:8D 25<br>$\mu\text{mol m}^{-2} \text{s}^{-1}$ | | D:D | | L:L 5<br>$\mu\text{mol m}^{-2} \text{s}^{-1}$ | | L:L 17<br>$\mu\text{mol m}^{-2} \text{s}^{-1}$ | | L:L 25<br>$\mu\text{mol m}^{-2} \text{s}^{-1}$ | | 16L:8D 75<br>$\mu\text{mol m}^{-2} \text{s}^{-1}$ | | L:L 50<br>$\mu\text{mol m}^{-2} \text{s}^{-1}$ | | L:L 75<br>$\mu\text{mol m}^{-2} \text{s}^{-1}$ | | 6L:6D 25<br>$\mu\text{mol m}^{-2} \text{s}^{-1}$ | |
| --- | --- | --- | --- | --- | --- | --- | --- | --- | --- | --- | --- | --- | --- | --- | --- | --- | --- | --- |
| rep. number | 3 |  | 4 |  | 3 |  | 10 |  | 7 |  | 4 |  | 4 |  | 8 |  | 4 |  |
| ERP rhythmic lines | 3/3 |  | 0/4 |  | 0/3 |  | 9/10 |  | 7/7 |  | 4/4 |  | 4/4 |  | 6/8 |  | 4/4 |  |
| FFT-NLLS algorithm | mean | s.d. | mean | s.d. | mean | s.d. | mean | s.d. | mean | s.d. | mean | s.d. | mean | s.d. | mean | s.d. | mean | s.d. |
| Period | 23,98 | 0,23 | N. A. | N. A. | N. A. | N. A. | 27,77 | 1,19 | 28,54 | 1,64 | 23,84 | 0,99 | 27,84 | 3,96 | 25,98 | 2,42 | 11,78 | 0,16 |
| Phase | 13,28 | 0,54 | N. A. | N. A. | N. A. | N. A. | 12,7 | 1,04 | 11,62 | 0,79 | 14,33 | 1,45 | 13,01 | 2,3 | 14,02 | 2,07 | 14,40 | 1,50 |
| Amplitude | 1,41 | 0,04 | N. A. | N. A. | N. A. | N. A. | 0,63 | 0,19 | 0,43 | 0,03 | 0,12 | 0,01 | 0,28 | 0,03 | 0,29 | 0,05 | 0,90 | 0,15 |
| RAE | 0,16 | 0,03 | N. A. | N. A. | N. A. | N. A. | 0,43 | 0,11 | 0,42 | 0,1 | 0,25 | 0,07 | 0,46 | 0,11 | 0,56 | 0,11 | 0,64 | 0,08 |
| div. per day | 0,97 | 0,03 | N. A. | N. A. | 0,20 | 0,07 | 1,07 | 0,19 | 1,09 | 0,06 | 1,49 | 0,09 | 1,24 | 0,05 | 1,4 | 0,03 | 0,49 | 0,14 |

Table S2: Rhythmicity parameters and growth of *P. tricornutum* cells grown under different L:D cycles of different light intensities and periods and following a shift to different free running conditions.
| Free running Temperature | 14°C |  | 16°C |  | 18°C |  | 20°C |  | 22°C |  |
| --- | --- | --- | --- | --- | --- | --- | --- | --- | --- | --- |
| rep. number | 8 |  | 3 |  | 10 |  | 3 |  | 8 |  |
| ERP rhythmic lines | 8/8 |  | 3/3 |  | 9/10 |  | 3/3 |  | 3/8 |  |
| FFT-NLLS algorithm | mean | s.d. | mean | s.d. | mean | s.d. | mean | s.d. | mean | s.d. |
| Period | 24,70 | 1,85 | 23,84 | 2,77 | 27,77 | 1,19 | 27,28 | 0,93 | 25,67 | 4,37 |
| Phase | 14,34 | 3,04 | 9,37 | 5,72 | 12,7 | 1,04 | 11,87 | 1,43 | 16,62 | 2,93 |
| Amplitude | 0,58 | 0,10 | 0,84 | 0,27 | 0,63 | 0,19 | 0,72 | 0,19 | 0,88 | 0,29 |
| RAE | 0,50 | 0,14 | 0,46 | 0,23 | 0,43 | 0,11 | 0,53 | 0,20 | 0,66 | 0,05 |
| div. per day | 0,97 | 0,10 | 1,22 | 0,03 | 1,18 | 0,03 | 1,33 | 0,06 | 1,05 | 0,13 |

**Table S3:** Rhythmicity parameters and growth of *P. tricornutum* WT, CTR, *RITMO1* KO1, KO2 and OE1 adapted to different photoperiods. The experiments were realised in cells adapted to 25 µmol photons m^−2^ s^−1^ at different photoperiods as represented in Fig. 2: 16L:8D; 12L:12D; 8L:16D. Rep. number indicates the number of biological replicas used for the analyses. EPR rhythmic lines indicates the lines that passed the EPR rhythmicity test for each condition and that are used for the FFT-NLLS rhythmicity test below. The FFT-NLLS was used to get the period, phase, amplitude, the Relative Amplitude of Error (RAE) that represents the amplitude of the error between the fit used by the algorithm and the data divided by the amplitude of the oscillations. The cell division/day was calculated in cells in exponential phase over the course of four days (n=3) for all lines. Statistical differences were examined using unpaired Student’s t-test with the WT as reference sample. Data for 16L:8D are the same than those shown in Table S1, here reproposed to facilitate the comparisons between the different conditions.

| 16L:8D | WT |  | CTR |  |  | OE1 |  |  | KO1 |  |  | KO2 |  |  |
| --- | --- | --- | --- | --- | --- | --- | --- | --- | --- | --- | --- | --- | --- | --- |
| rep. Number | 3 |  | 3 |  |  | 3 |  |  | 3 |  |  | 3 |  |  |
| ERP rhythmic lines | 3/3 |  | 3/3 |  |  | 3/3 |  |  | 3/3 |  |  | 3/3 |  |  |
| FFT-NLLS algorithm | mean | s.d. | mean | s.d. | p. value | mean | s.d. | p. value | mean | s.d. | p. value | mean | s.d. | p. value |
| Period | 23,98 | 0,23 | 24,25 | 0,26 | n. s. | 23,09 | 0,19 | 0,0067 | 24,00 | 0,15 | n. s. | 24,21 | 0,11 | n. s. |
| Phase | 13,28 | 0,54 | 12,16 | 0,52 | n. s. | 14,38 | 0,60 | n. s. | 13,03 | 0,37 | n. s. | 12,44 | 0,41 | n. s. |
| Amplitude | 1,41 | 0,04 | 1,31 | 0,28 | n. s. | 1,06 | 0,05 | 0,0007 | 1,32 | 0,17 | n. s. | 1,34 | 0,10 | n. s. |
| RAE | 0,16 | 0,01 | 0,16 | 0,04 | n. s. | 0,33 | 0,05 | 0,0095 | 0,19 | 0,03 | n. s. | 0,20 | 0,02 | n. s. |
| div. per day | 1,15 | 0,03 | 1,09 | 0,02 | n. s. | 1,06 | 0,09 | n. s. | 0,91 | 0,17 | n. s. | 0,90 | 0,15 | n. s. |

| 12L:12D | WT |  | CTR |  |  | OE1 |  |  | KO1 |  |  | KO2 |  |  |
| --- | --- | --- | --- | --- | --- | --- | --- | --- | --- | --- | --- | --- | --- | --- |
| rep. Number | 4 |  | 3 |  |  | 3 |  |  | 3 |  |  | 3 |  |  |
| ERP rhythmic lines | 4/4 |  | 3/3 |  |  | 3/3 |  |  | 3/3 |  |  | 3/3 |  |  |
| FFT-NLLS algorithm | mean | s.d. | mean | s.d. | p. value | mean | s.d. | p. value | mean | s.d. | p. value | mean | s.d. | p. value |
| Period | 24,47 | 0,38 | 23,84 | 0,28 | n. s. | 24,44 | 0,58 | n. s. | 24,13 | 0,27 | n. s. | 25,08 | 1,04 | n. s. |
| Phase | 10,65 | 1,00 | 11,85 | 0,14 | n. s. | 10,88 | 1,23 | n. s. | 11,28 | 0,90 | n. s. | 10,23 | 1,52 | n. s. |
| Amplitude | 1,12 | 0,22 | 1,08 | 0,13 | n. s. | 0,91 | 0,13 | n. s. | 1,05 | 0,09 | n. s. | 1,15 | 0,06 | n. s. |
| RAE | 0,15 | 0,03 | 0,12 | 0,01 | n. s. | 0,14 | 0,02 | n. s. | 0,15 | 0,05 | n. s. | 0,19 | 0,03 | n. s. |
| div. per day | 0,81 | 0,03 | 0,72 | 0,06 | n. s. | 0,70 | 0,06 | n. s. | 0,65 | 0,06 | n. s. | 0,75 | 0,03 | n. s. |

Table S3: Rhythmicity parameters and growth of *P. tricornutum* WT, CTR, *RITMO1* KO1, KO2 and OE1 adapted to different photoperiods.
| 8L:16D | WT |  | CTR |  |  | OE1 |  |  | KO1 |  |  | KO2 |  |  |
| --- | --- | --- | --- | --- | --- | --- | --- | --- | --- | --- | --- | --- | --- | --- |
| rep. Number | 4 |  | 3 |  |  | 3 |  |  | 3 |  |  | 3 |  |  |
| ERP rhythmic lines | 4/4 |  | 3/3 |  |  | 3/3 |  |  | 3/3 |  |  | 3/3 |  |  |
| FFT-NLLS algorithm | mean | s.d. | mean | s.d. | p. value | mean | s.d. | p. value | mean | s.d. | p. value | mean | s.d. | p. value |
| Period | 23,63 | 0,16 | 23,83 | 0,14 | n. s. | 24,36 | 0,55 | 0,0486 | 24,05 | 0,46 | n. s. | 24,68 | 0,99 | n. s. |
| Phase | 9,78 | 0,13 | 9,84 | 0,44 | n. s. | 9,00 | 0,70 | n. s. | 9,73 | 0,23 | n. s. | 8,58 | 1,70 | n. s. |
| Amplitude | 0,27 | 0,03 | 0,24 | 0,03 | n. s. | 0,22 | 0,02 | 0,0332 | 0,25 | 0,03 | n. s. | 0,22 | 0,04 | n. s. |
| RAE | 0,19 | 0,03 | 0,23 | 0,09 | n. s. | 0,19 | 0,06 | n. s. | 0,21 | 0,07 | n. s. | 0,27 | 0,01 | 0,0090 |
| div. per day | 0,79 | 0,04 | 0,78 | 0,01 | n. s. | 0,85 | 0,03 | n. s. | 0,76 | 0,02 | n. s. | 0,76 | 0,04 | n. s. |

**Table S4:** Rhythmicity parameters and growth of *P. tricornutum* WT, CTR, *RITMO1* OE1, KO1, KO2 and KO2 complemented line KO2-C2 in free-running L:L conditions. The experiments were realised with *P. tricornutum* WT, *RITMO1* KO1, KO2, OE1 and KO2-C2 strains adapted to 16L:8D 25 µmol photons m^−2^ s^−1^ and transferred to L:L of 17 µmol photons m^−2^ s^−1^ (Fig. 3 and Fig. 7). Rep. number indicates the number of biological replicas used for the analysis. EPR rhythmic lines indicates the lines that passed the EPR rhythmicity test for each condition and that are used for the FFT-NLLS rhythmicity test below. The FFT-NLLS was used to get the period, phase, amplitude, the Relative Amplitude of Error (RAE) that represents the amplitude of the error between the fit used by the algorithm and the data divided by the amplitude of the oscillations. The cell division/day was calculated in cells in exponential phase over the course of four days (n=6) for all lines. Statistical differences were examined using unpaired Student’s t-test with the WT as reference sample.

|  | WT |  | CTR |  |  | OE1 |  |  | KO1 |  |  | KO2 |  |  | KO2-C2 |  |  |
| --- | --- | --- | --- | --- | --- | --- | --- | --- | --- | --- | --- | --- | --- | --- | --- | --- | --- |
| rep. number | 17 |  | 12 |  |  | 7 |  |  | 17 |  |  | 15 |  |  | 8 |  |  |
| ERP rhythmic lines | 17/17 |  | 12/12 |  |  | 5/7 |  |  | 11/17 |  |  | 9/15 |  |  | 7/8 |  |  |
| FFT-NLLS algorithm | mean | s.d. | mean | s.d. | p. value | mean | s.d. | p. value | mean | s.d. | p. value | mean | s.d. | p. value | mean | s.d. | p. value |
| Period | 28,58 | 2,27 | 28,80 | 1,53 | n.s. | 28,12 | 7,57 | n.s. | 25,83 | 2,69 | 0,0072 | 24,93 | 6,72 | 0,0008 | 26,89 | 3,13 | n.s. |
| Phase | 12,16 | 1,36 | 12,62 | 1,32 | n.s. | 12,48 | 2,38 | n.s. | 14,05 | 2,19 | 0,0091 | 13,73 | 2,58 | n.s. | 13,25 | 4,92 | n.s. |
| Amplitude | 0,96 | 0,45 | 0,97 | 0,50 | n.s. | 0,62 | 0,39 | n.s. | 0,89 | 0,42 | n.s. | 0,87 | 0,54 | n.s. | 0,99 | 0,29 | n.s. |
| RAE | 0,39 | 0,16 | 0,37 | 0,20 | 0,6739 | 0,69 | 0,11 | 0,0001 | 0,58 | 0,18 | 0,0015 | 0,63 | 0,14 | 0,0008 | 0,45 | 0,16 | n.s. |
| div. per day | 1,18 | 0,13 | 1,26 | 0,11 | n.s. | 1,23 | 0,11 | n.s. | 1,18 | 0,15 | n.s. | 1,23 | 0,19 | n.s. | 1,23 | 0,11 | n.s. |

**Table S5:** Analysis of photosynthetic parameters rhythmicity in *P. tricornutum* WT, *RITMO1* KO1, KO2 and OE1 in L:D and L:L. The experiments were performed with *P. tricornutum* WT, *RITMO1* KO1, KO2 and OE1 strains grown under 16L:8D (50 µmol photons m^−2^ s^−1^) and in L:L (30 µmol photons m^−2^ s^−1^) as described in Fig. 5. JTK cycle analysis was performed on the dataset obtained from four biological replicas for each strain with the replica option selected (n=4). In L:D, due to the reduced period of sampling a 24 h period was enforced. In L:L, the period was free to vary between 20 and 28 h. Rhythmicity p. value indicates the p. value of rhythmicity provided by JTK cycle algorithm after a Benjamini-Hochberg correction. Phase design the acrophase relative to the first L/D or theoric L/D transition. Amp. designs the amplitude of the variation. Rhythmicity p. value significance is noted with stars, * = p<0.05; ** = p<0.01;*** = p<0.001.

| Parameter | 16L:8D |  |  | L:L |  |  |  |
| --- | --- | --- | --- | --- | --- | --- | --- |
|  | Rhythmicity p. value | Phase | Amp. | Rhythmicity p. value | Period | Phase | Amp. |
| FvFm KO1 | 5,13E-05 *** | 8 | 0,041 | 1,00E+00 | 28 | 22 | 0,008 |
| FvFm KO2 | 1,05E-05 *** | 8 | 0,043 | 1,00E+00 | 24 | 16 | 0,004 |
| FvFm OE1 | 4,17E-08 *** | 8 | 0,04 | 1,66E-01 | 24 | 16 | 0,007 |
| FvFm WT | 1,31E-06 *** | 8 | 0,044 | 9,00E-07 *** | 24 | 16 | 0,013 |
| E50NPQ KO1 | 1,35E-02 * | 20 | 31,9 | 1,00E+00 | 28 | 4 | 9,13 |
| E50NPQ KO2 | 3,25E-04 *** | 18 | 36,2 | 1,00E+00 | 20 | 18 | 6,44 |
| E50NPQ OE1 | 1,05E-05 *** | 18 | 40 | 1,00E+00 | 20 | 22 | 8,74 |
| E50NPQ WT | 5,69E-06 *** | 18 | 42,3 | 6,08E-05 *** | 28 | 2 | 28,2 |
| NPQrel KO1 | 2,76E-04 *** | 20 | 20,4 | 1,00E+00 | 28 | 6 | 2,9 |
| NPQrel KO2 | 1,01E-03 ** | 18 | 23,7 | 9,47E-02 | 28 | 6 | 2,66 |
| NPQrel OE1 | 4,18E-08 *** | 16 | 22,6 | 1,00E+00 | 28 | 22 | 1,85 |
| NPQrel WT | 9,77E-05 *** | 18 | 27,4 | 2,95E-07 *** | 24 | 8 | 11,7 |
| σPSII KO1 | 4,91E-07 *** | 22 | 16,8 | 1,00E+00 | 28 | 4 | 1,84 |
| σPSII KO2 | 4,26E-05 *** | 22 | 21,2 | 1,00E+00 | 20 | 6 | 3,82 |
| σPSII OE1 | 5,07E-06 *** | 20 | 14 | 1,66E-01 | 24 | 4 | 3,18 |
| σPSII WT | 5,69E-06 *** | 22 | 18 | 8,66E-03 ** | 24 | 4 | 5,37 |
| rETRm KO1 | 1,57E-06 *** | 8 | 7,76 | 1,00E+00 | 28 | 6 | 0,803 |
| rETRm KO2 | 4,18E-06 *** | 8 | 7,52 | 1,00E+00 | 28 | 10 | 1,62 |
| rETRm OE1 | 2,13E-07 *** | 8 | 6,65 | 1,00E+00 | 28 | 14 | 1,19 |
| rETRm WT | 9,32E-09 *** | 8 | 7,96 | 1,00E+00 | 28 | 10 | 0,818 |
| alpha KO1 | 2,22E-02 * | 8 | 0,044 | 4,46E-01 | 28 | 22 | 0,036 |
| alpha KO2 | 1,04E-02 * | 12 | 0,047 | 1,00E+00 | 28 | 24 | 0,032 |
| alpha OE1 | 2,00E-02 * | 12 | 0,053 | 9,70E-01 | 28 | 26 | 0,012 |
| alpha WT | 6,08E-02 | 8 | 0,043 | 1,00E+00 | 20 | 14 | 0,006 |
| NPQm KO1 | 2,58E-06 *** | 6 | 0,162 | 1,00E+00 | 28 | 8 | 0,078 |
| NPQm KO2 | 9,16E-06 *** | 6 | 0,157 | 1,00E+00 | 28 | 8 | 0,07 |
| NPQm OE1 | 1,11E-06 *** | 4 | 0,144 | 9,64E-02 | 28 | 12 | 0,062 |
| NPQm WT | 2,77E-05 *** | 6 | 0,14 | 1,00E+00 | 28 | 10 | 0,047 |

**Fig. S1:**
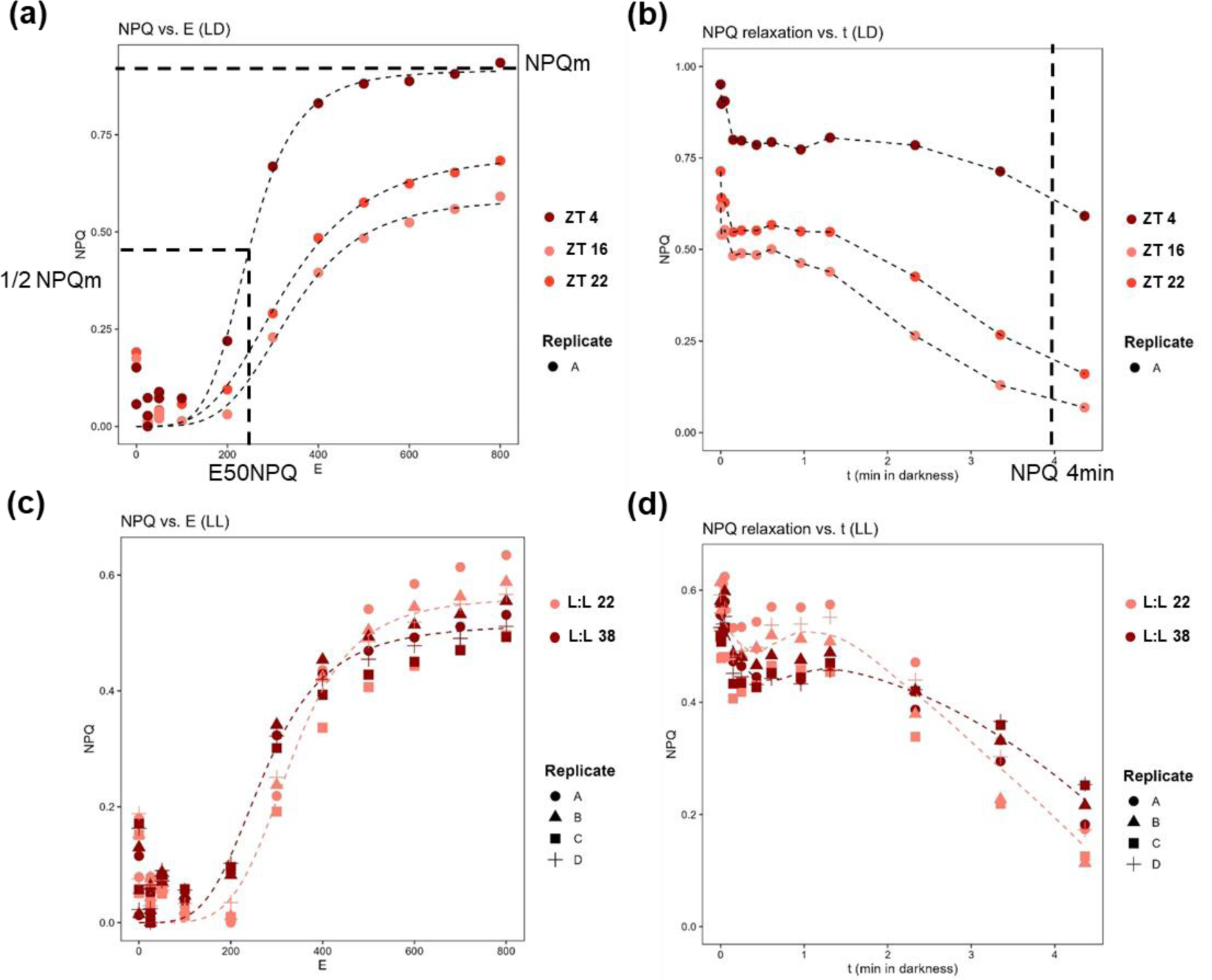
Evolution of non-photochemical quenching (NPQ) versus light intensity and kinetics of NPQ relaxation in darkness. (a, c) Relation between non-photochemical quenching (NPQ) and light intensity (E) for *P. tricornutum* WT strain of under 16L:8D 25 µmol photons m^−2^ s^−1^ (a) and transferred to L:L 17 µmol photons m^−2^ s^−1^ (c) at various sampling time. Light intensity for which 50% of maximum NPQ (E50NPQ) and maximum NPQ (NPQm) are indicated by thick dashed lines. (b, d) Relaxation of the NPQ in the dark after NPQm induction for WT strain (Pt1) replicate A in both L:D (b) and L:L (d) at various sampling times, in L:L all replicates around L:L 2 and L:L 18 are shown to display maximal contrast in the parameters values. Light dashed lines represent the fit for this parameter (Materials and Methods). The NPQ 4 min used to calculate NPQrel (%age of maximal NPQ monitored relaxed after 4 min in dark) is indicated by thick dashed lines.

**Fig. S2:**
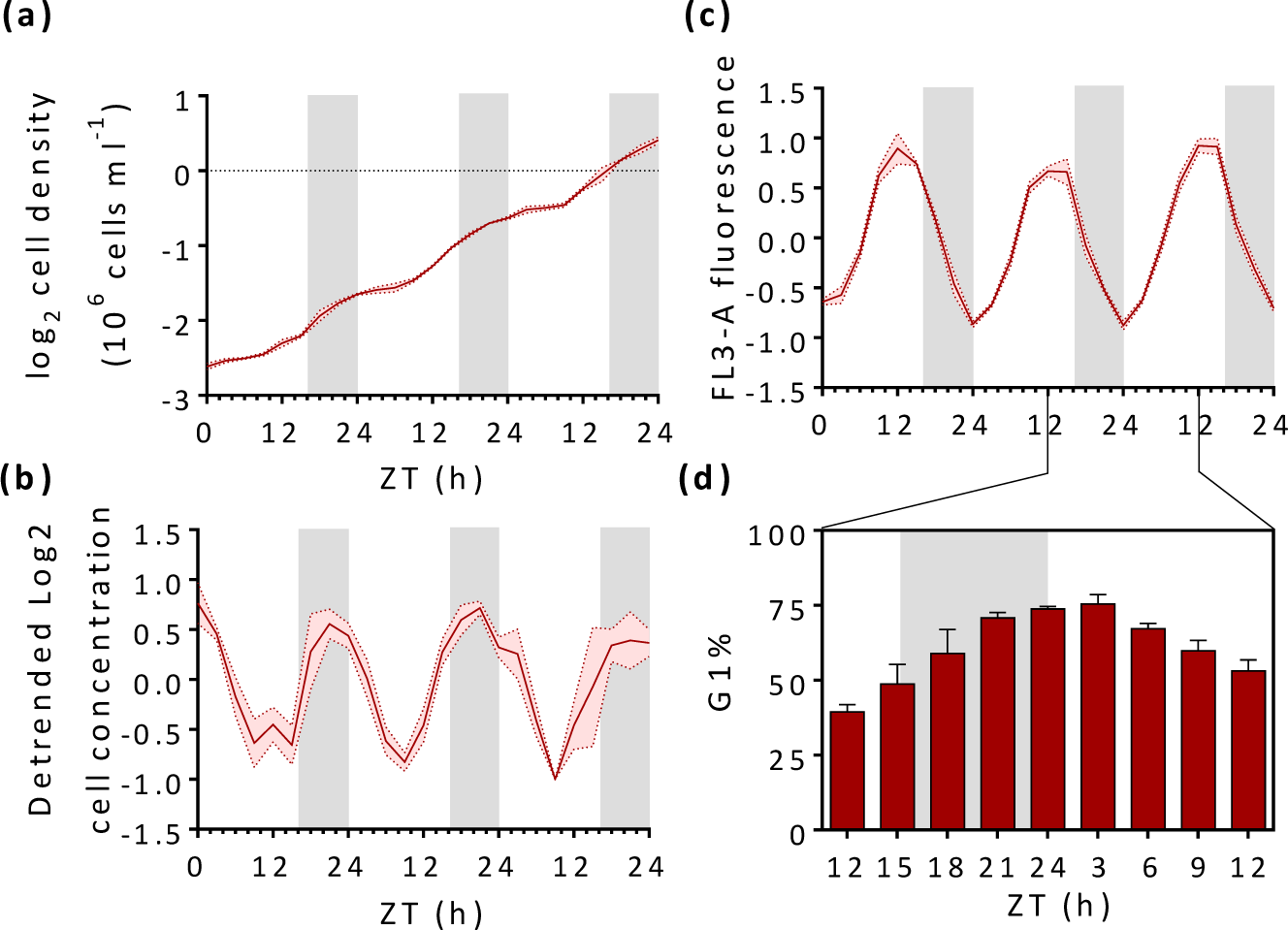
Characterisation of growth (a, b), cellular fluorescence (c) and cell cycle synchronization (d) in *P. tricornutum* cells (n>=3) entrained under 16L:8D 25 µmol photons m^−2^ s^−1^. Samples were collected every 3 h for the analyses. Cell concentration plotted as 10^6^ cells ml^−1^ (a). Log2 of the cell concentration presented in panel a after a baseline detrending to highlight diel variations (b). Cellular fluorescence (FL3-A parameter) profile of the data reported in panel a and b (c). Solid lines represent the average, coloured dashed lines represent SD. Percentage of cells in G1 phase in the highlighted time window (d). Cell phase was determined by flow cytometry after DAPI staining of the cellular DNA content (V1-A parameter). Red bars represent the average and error bar the SD. White and grey regions represent light and dark periods.

**Fig. S3.**
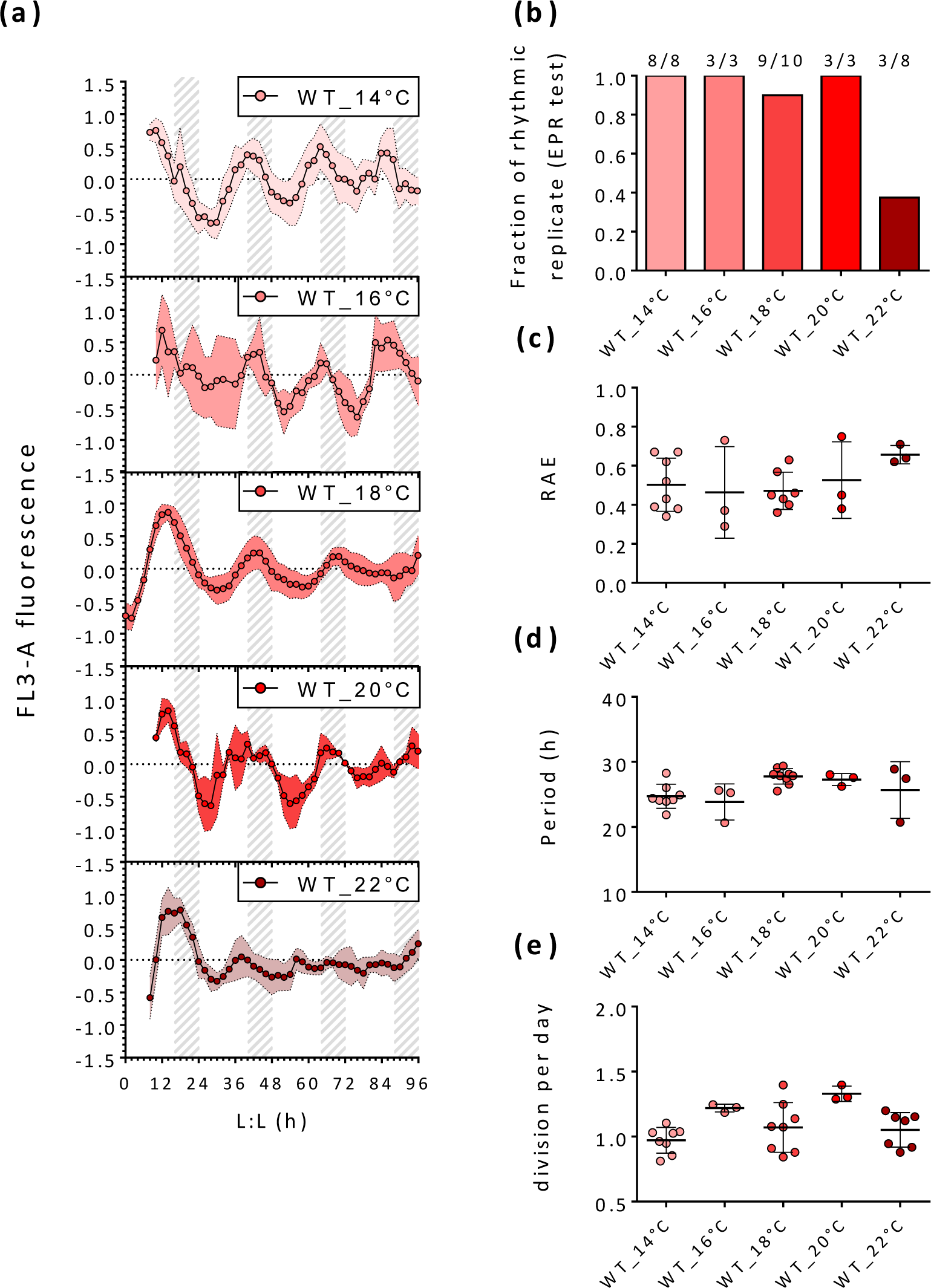
Characterization of cellular fluorescence rhythmicity in *P. tricornutum* WT strain entrained under 16L:8D 25 µmol photons m^−2^ s^−1^ at 18°C and after the switch to L:L 17 µmol photons m^−2^ s^−1^ at various temperatures. Shift to lower temperature were done at the beginning of the last night (ZT 16) and shift to upper temperature at the end of the last night (ZT 24). Normalised and baseline detrended circadian cellular FL3-A fluorescence profiles in WT at 14°C, 16°C, 18°C, 20°C and 22°C (a) (n>=3). Dots represent mean FL3-A fluorescence values, coloured dashed lines represent SD. Grey dashed regions represent subjective nights in free-run conditions. (b) Fraction of replicates that passes EPR test for all conditions. (c) Relative amplitude of Error (RAE) of the FFT-NLLS method fit for the lines found rhythmic with EPR test. (d) Period estimation obtained with FFT-NLLS method. (e) Division per day along the experiment (n>=3). Dots represent mean values, error bars represent SD.

**Fig. S4:**
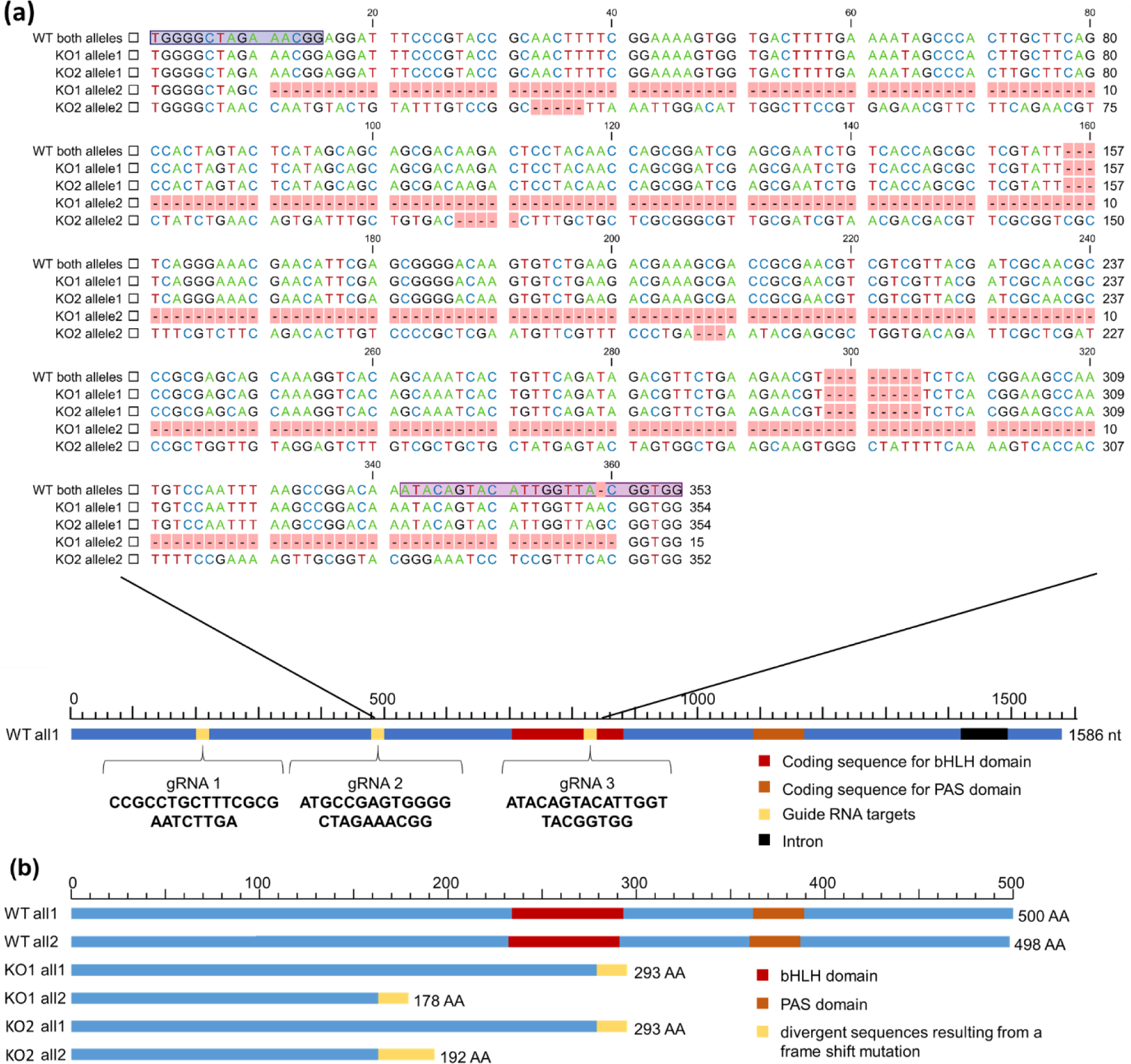
Analysis of *RITMO1* gene modification in knock-out (KO) lines compared to the wild-type (WT) sequence. (a) Schematic representation of the *P. tricornutum RITMO1* locus on Chromosome 5 (Phatr3_J44962 in https://www.diatomicsbase.bio.ens.psl.eu/). Intron is represented with a black box. Red and orange boxes represent the relative location of the coding sequence for bHLH and PAS domains, respectively. Yellow boxes highlight the localisation of the target sequences (underlined) of the three gRNAs used for *RITMO1* CRISPR-Cas9 mutagenesis. On the top is shown the alignment of the regions containing the mutations in both alleles of *RITMO1* KOs compared to the WT sequence. b) Schematic representation of the RITMO1 protein encoded by the two alleles of the *P. tricornutum* WT strain and the two KO mutants used in this study and identified by Sanger sequencing. Red and orange boxes represent the relative location of bHLH and PAS domains respectively predicted using InterProScan. Yellow boxes highlight sequence divergence from wild type resulting from a frame shift mutation.

**Fig. S5.**
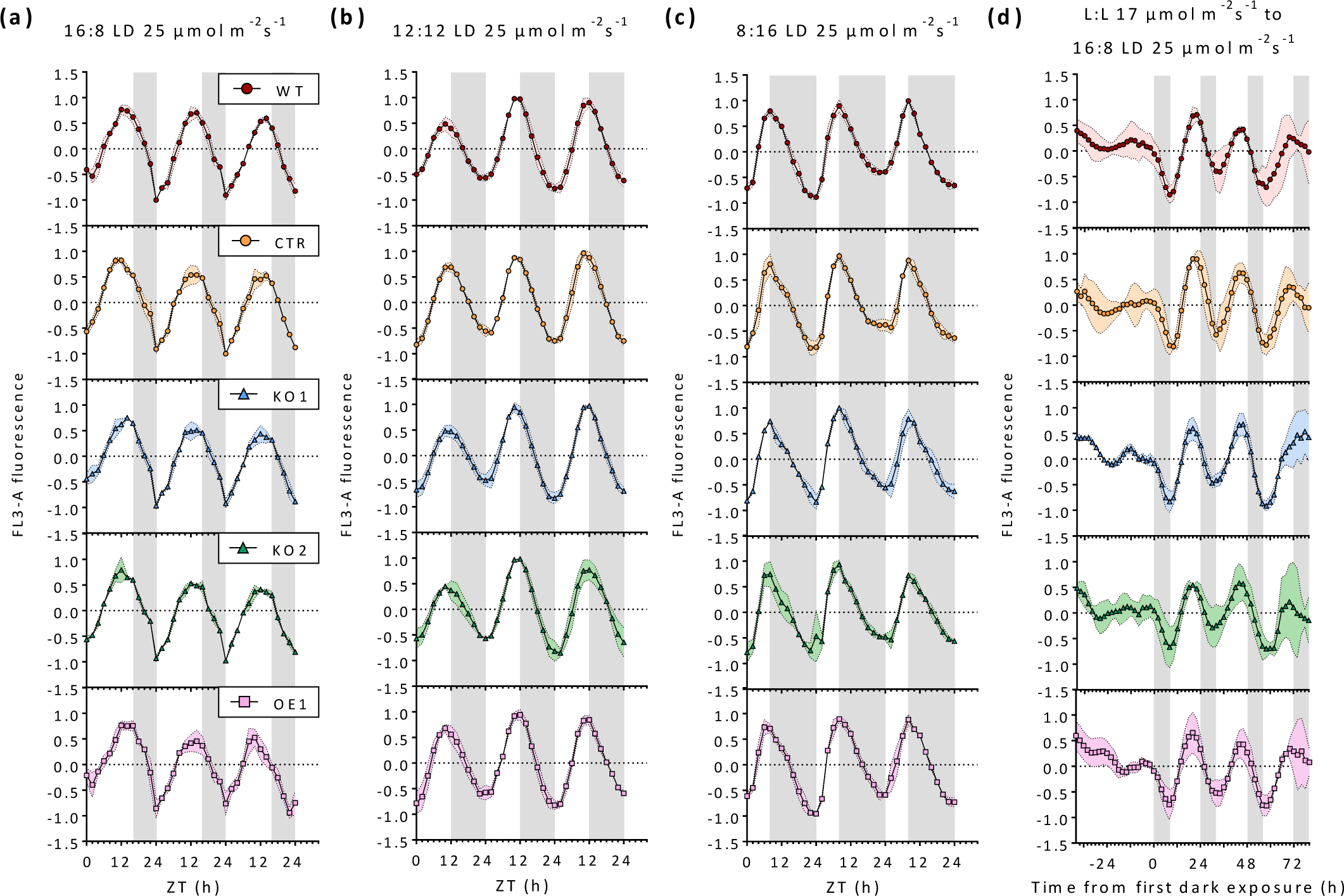
Characterization of cellular fluorescence rhythmicity of *P. tricornutum* WT, CTR, and *RITMO1* OE1, KO1 and KO2 lines (n>=3) under L:D cycles. In (a-c), the experimental conditions and data are the same as in Fig. 2, but are represented as individual data for each strain to improve visualization of results. (d), re-entrainment experiments as for Fig. 4, with the same criteria.

**Fig. S6:**
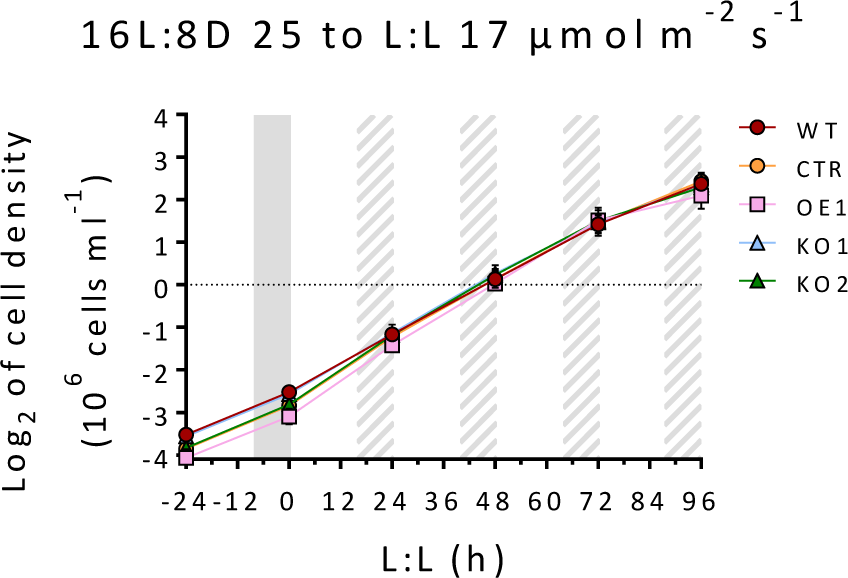
Analysis of growth capacity in *P. tricornutum* WT, CTR, *RITMO1* KO1, KO2 and OE1. Growth curve of *P. tricornutum* WT, CTR, *RITMO1* KO1, KO2 and OE1 cells (n=3) under 16L:8D 25 µmol photons m^−2^ s^−1^ and transferred to L:L 17 µmol photons m^−2^ s^−1^. Dots represent mean value of the log_2_ of the cell density, error bar indicates SD.

**Fig. S7:**
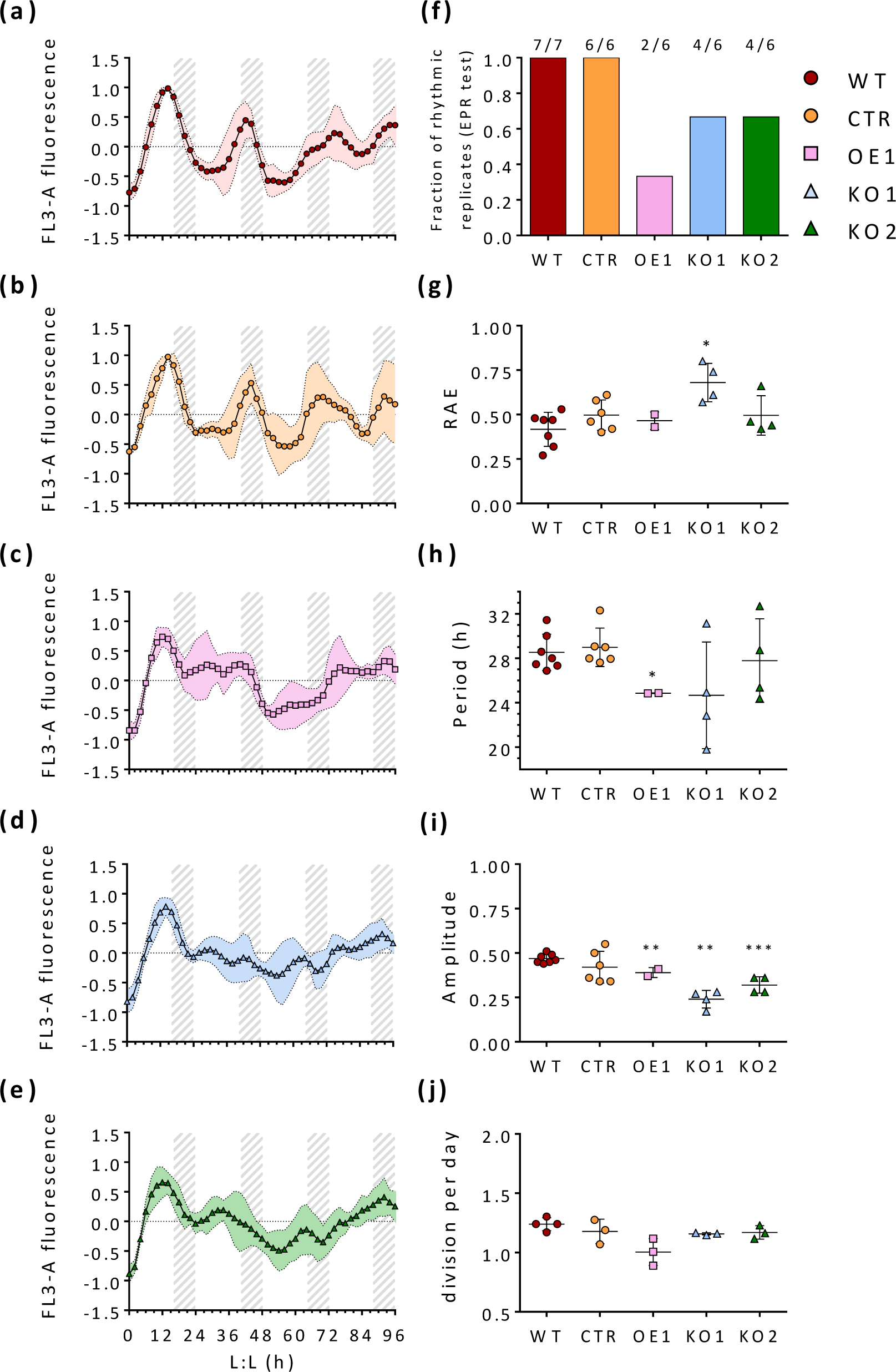
Characterization of cellular fluorescence rhythmicity in *P. tricornutum* WT, *RITMO1* KO1, KO2 and OE1 strains entrained at 16L:8D 25 µmol photons m^−2^ s^−1^ after the switch to L:L 25 µmol photons m^−2^ s^−1^. Normalised and baseline detrended circadian cellular FL3-A fluorescence profiles in WT (a), CTR (b), OE1 (c), KO1 (d) and KO2 (e) (n>=6). Dots represent mean FL3-A fluorescence values, coloured dashed lines represent SD. Gray dashed regions represent subjective nights in free-run conditions. (f) Fraction of replicates that passes EPR test for all strains. (g) Relative amplitude of Error (RAE) of the FFT-NLLS method fit for the lines found rhythmic with EPR test. (h) Period estimation obtained with FFT-NLLS method. (i) Relative amplitude of Error of the FFT-NLLS method fit vs the predicted period for the lines found rhythmic with EPR test. (j) Division per day along the experiment (n>=3). Dots represent mean values, error bars represent SD. Statistical differences were examined using unpaired Student’s t-test with the WT as reference sample (* = p<0.05; ** = p<0.01;*** = p<0.001).

**Fig. S8:**
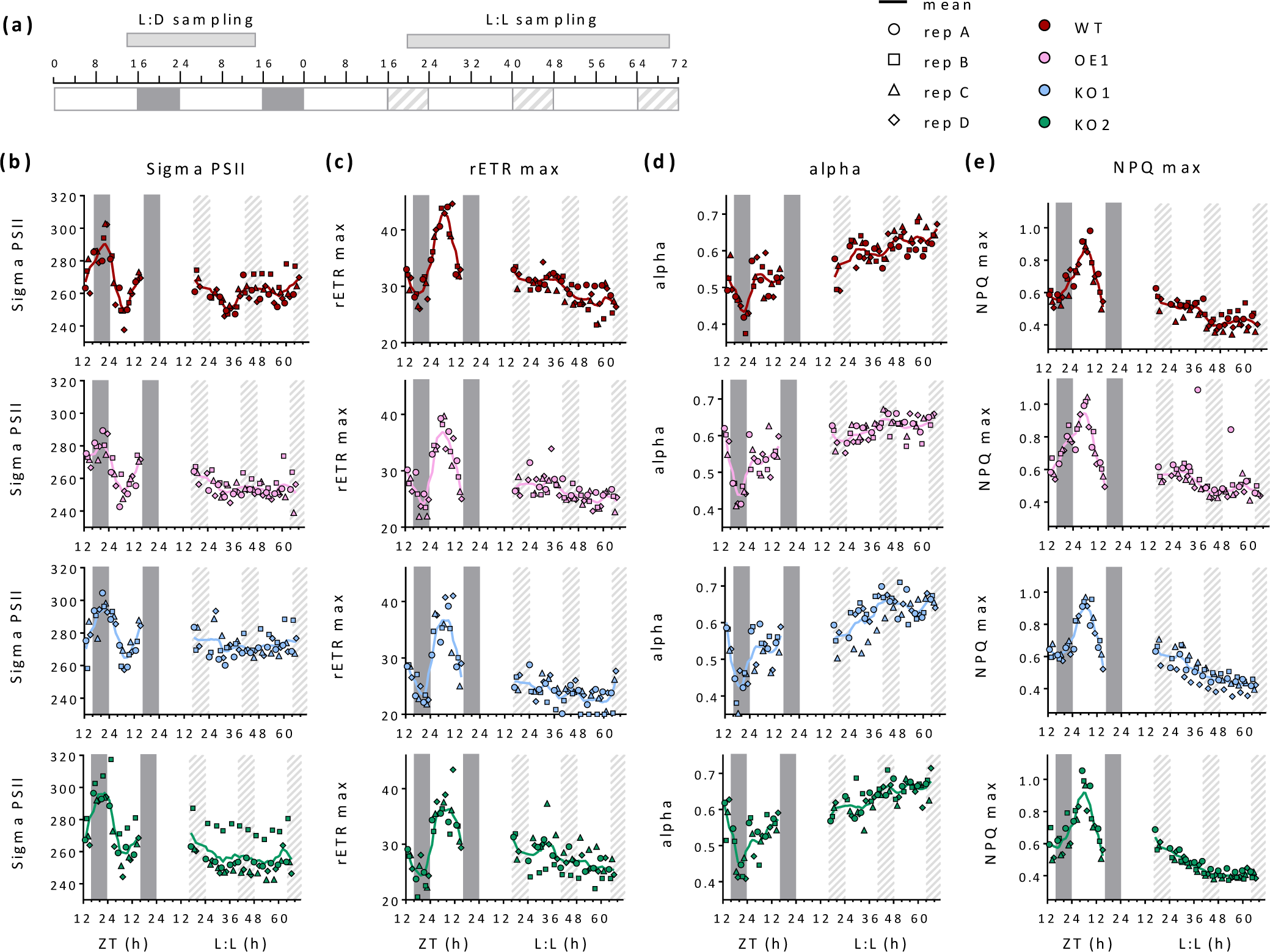
Analysis of photosynthetic parameters in *P. tricornutum* WT, *RITMO1* KO1, KO2 and OE1. (a) Schematic representation of sampling times for the analysis from WT, *RITMO1* KO1, KO2 and OE1 cells grown under 16L:8D 50 µmol photons m^−2^ s^−1^ and released to L:L 30 µmol photons m^−2^ s^−1^. Samples were collected continuously and measured one by one in the following order WT, KO1, KO2, OE1 from replicate A to replicate D, with 15 min interval due to the measurement. The measure of all strains and all replicates takes 4 h, measurement were performed over 24 h in L:D and 48 h in L:L. In free running, samples were collected starting at ZT16. (b) effective absorption cross-section of PSII (σPSII), (c) maximum of the relative electron transfer rate (rETR max), (d) initial light limited slope of rETR (alpha) and (e) maximal NPQ measured (NPQm). Dots, squares, triangles and diamonds represents the individual measures for biological replicates A to D, colored dashed lines represent the moving average of all replicates (window of 4 h, n=4). White and gray regions represent light and dark periods, gray dashed regions represent subjective nights in free-run conditions.

**Fig. S9:**
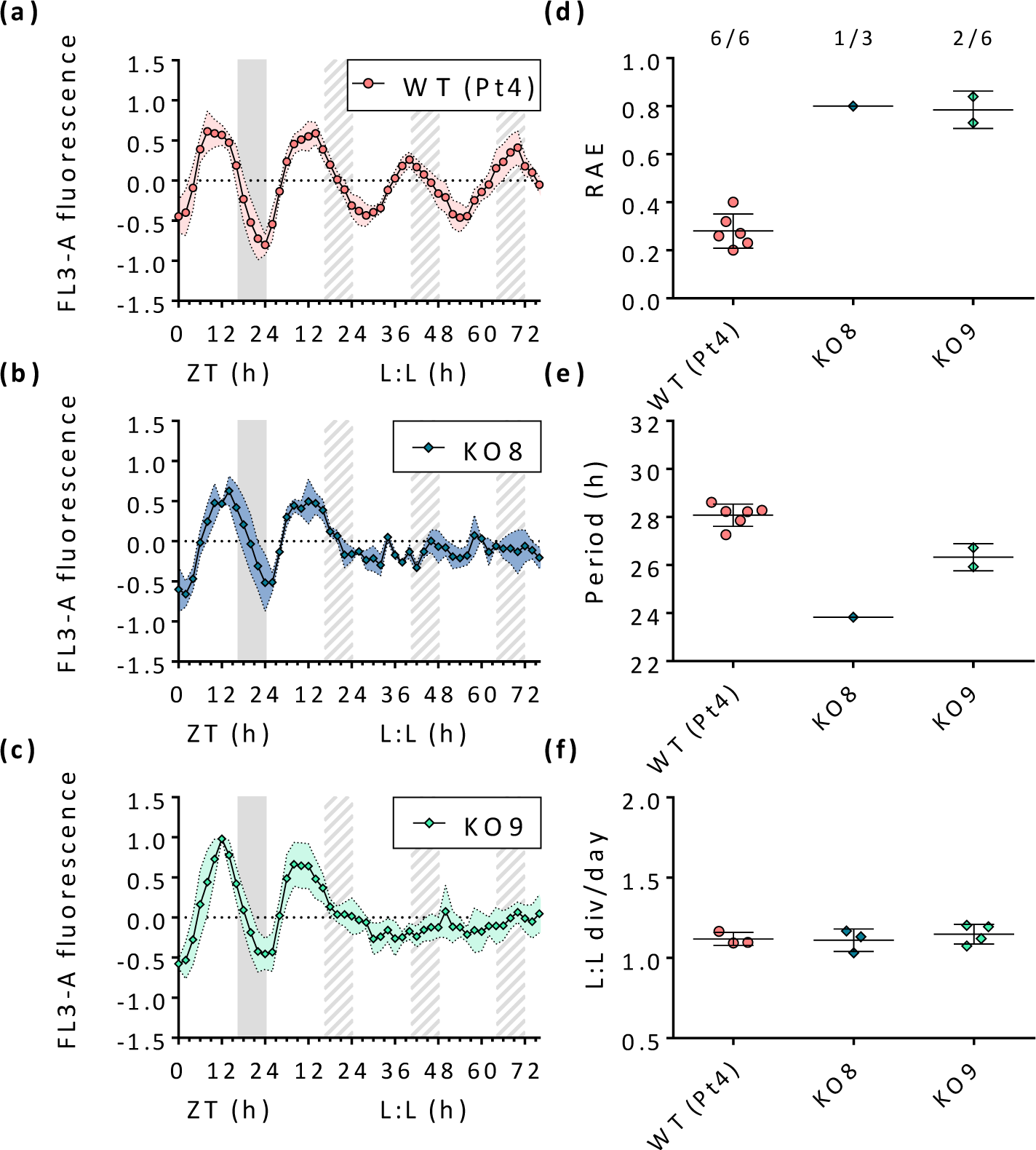
Characterization of cellular fluorescence rhythmicity in *P. tricornutum* WT (Pt4), *Aureo1a* KO8, KO9 strains under 16L:8D 25 µmol photons m^−2^ s^−1^ entrainment and following a switch to L:L 17 µmol photons m^−2^ s^−1^ for 3 days. Normalised and baseline detrended circadian cellular FL3-A fluorescence profiles in WT (Pt4) (a), Aureochrome1a KO8 (b) and KO9 (c) (n>=6). Dots represent mean FL3-A fluorescence values, coloured dashed lines represent SD. Gray dashed regions represent subjective nights in free-run conditions. (d) Relative amplitude of Error (RAE) of the FFT-NLLS method fit for the lines found rhythmic in L:L with EPR test, fraction of replicates that passes EPR test for all strains noted above. (e) Period estimation obtained with FFT-NLLS method. (f) Division rate along the experiment (n>=6).

**Fig. S10:**
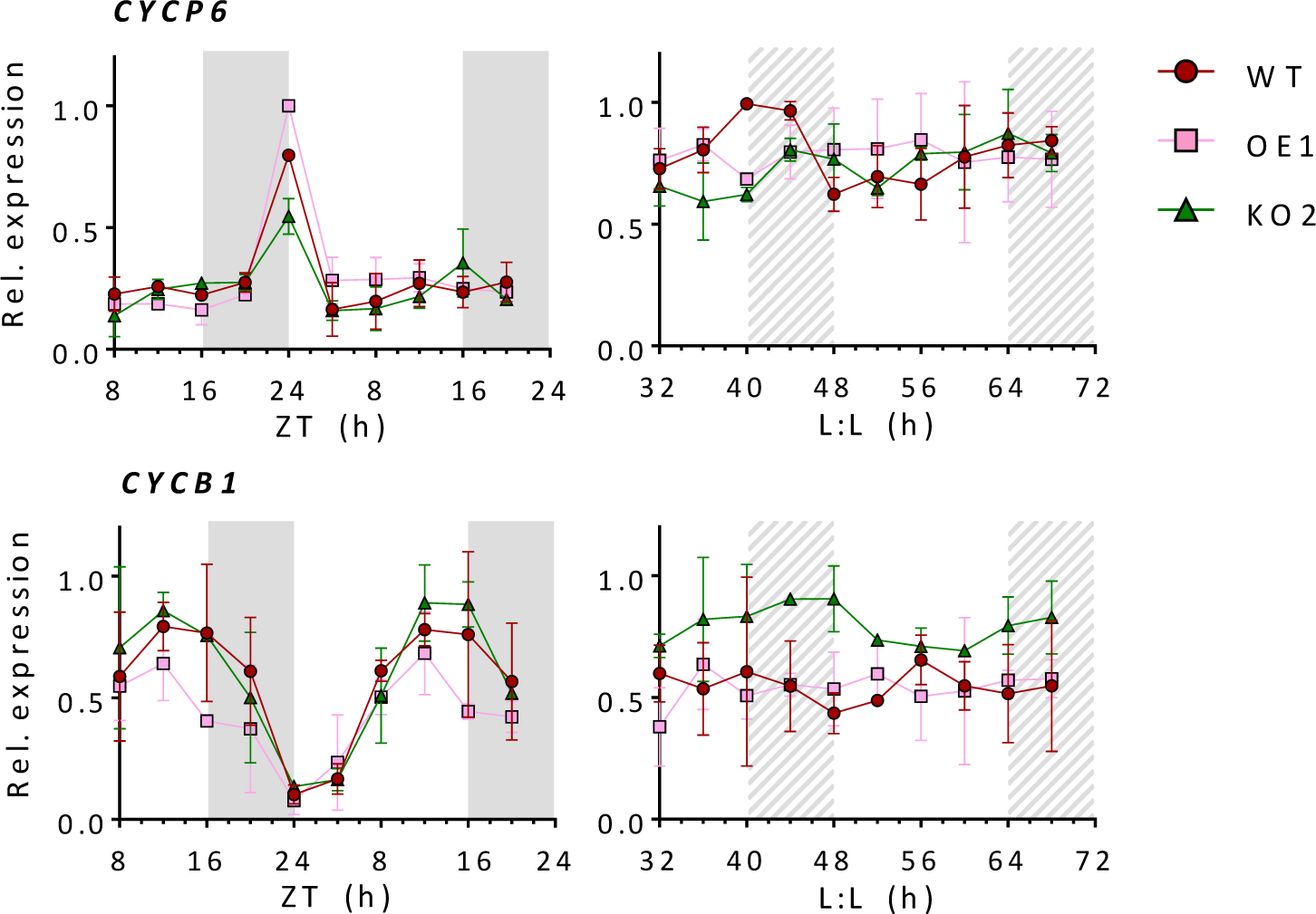
Analysis of the rhythmic expression of selected genes by qRT-PCR in *P. tricornutum* WT, *RITMO1* OE1 and KO2 lines under L:D cycles and L:L conditions. Expression profiles of the cyclin CYCP6 and CYCB1 for WT, KO2 and OE1 lines. Expression values represent the average of three biological triplicates ±SD are normalized using the *RPS*, *TBP* reference genes. Expression values are given relative to the maximum expression for each gene, where ‘1’ represents the highest expression value of the time series. Dots represent mean values, error bars represent SD (n=2). White and grey regions represent light and dark periods, grey dashed regions represent subjective nights in free-run conditions.

**Fig. S11:**
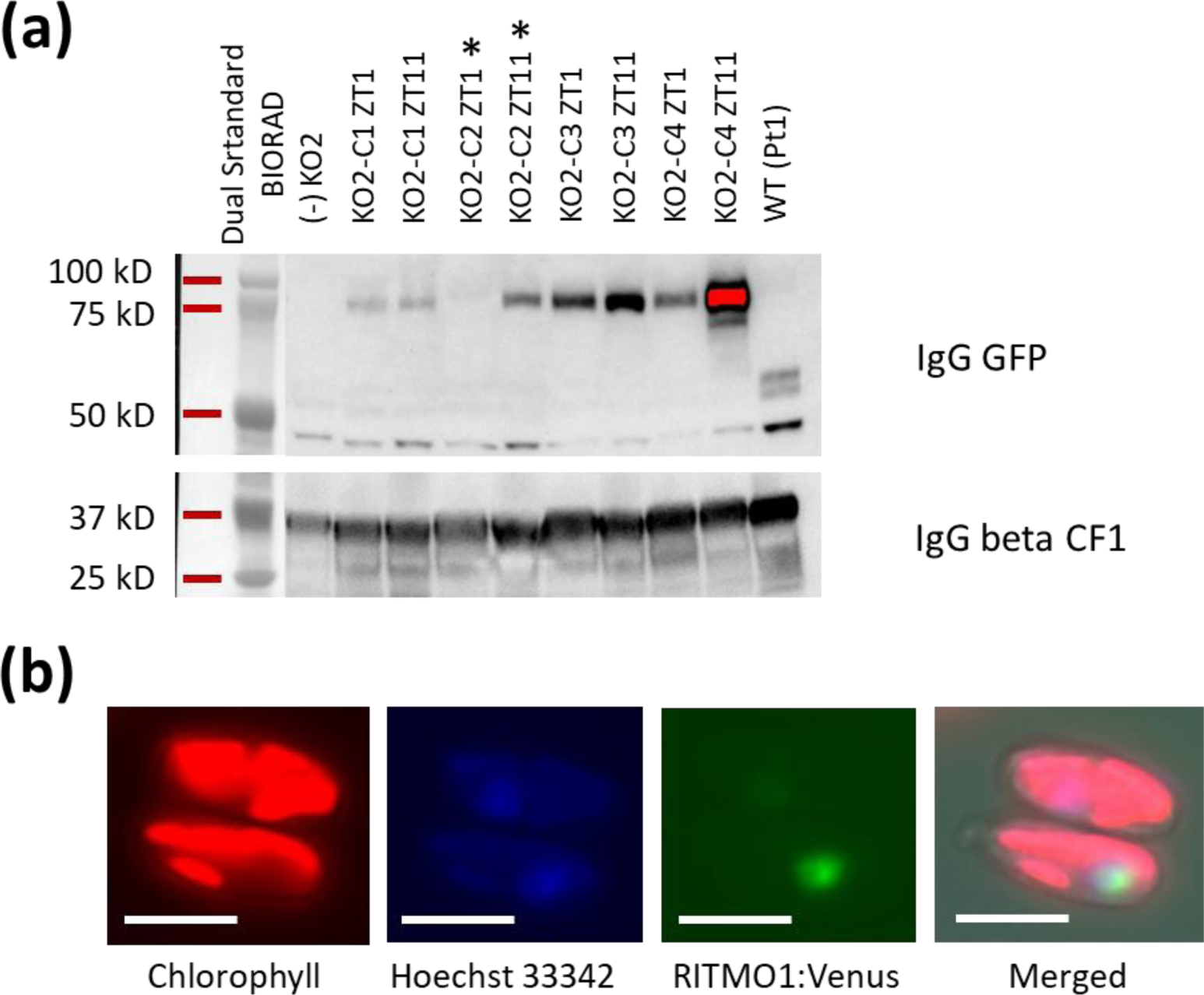
Selection of complemented *RITMO1* lines in *RITMO1* KO2 mutant. (a) Western blot analysis of transgenic lines of *RITMO1* KO2 mutant transformed with the RITMO1p:RITMO1g:Venus:RITMO1t construct by biolistic. In this experiment, cells were grown in 12L:12D cycles at 25 µmol photons m^−2^ s^−1^ and collected for the analysis 1h after light onset (ZT 1) and 1 h before the dark (ZT11). KO2 is the original mutant strain (negative control,-), KO2-C indicates the independent complemented lines. Top are the results with the IgG anti-GFP (1/2000) and down with an IgG anti-beta CF1 (1/20000) used as a loading control. The KO2-C2 line used in Fig. 7 is indicated with an asterisk. (b) Fluorescence microscopy of *P. tricornutum* cells expressing the RITMO1-Venus protein under the control of the endogenous promoter. Samples imaged on exponential phase cells grown in 16L:8D 25 µmol photons m^−2^ s^−1^ at ZT15. (Scale bar: 5 μm). For chlorophyll autofluorescence and Venus fluorescence cells were excited at 510 nm and detected at 650–741 nm and 529–562 nm, respectively. Nuclear DNA stained with Hoechst 33342 were visualized by illumination at 405 nm and detection at 424–462 nm.

**Fig. S12:**
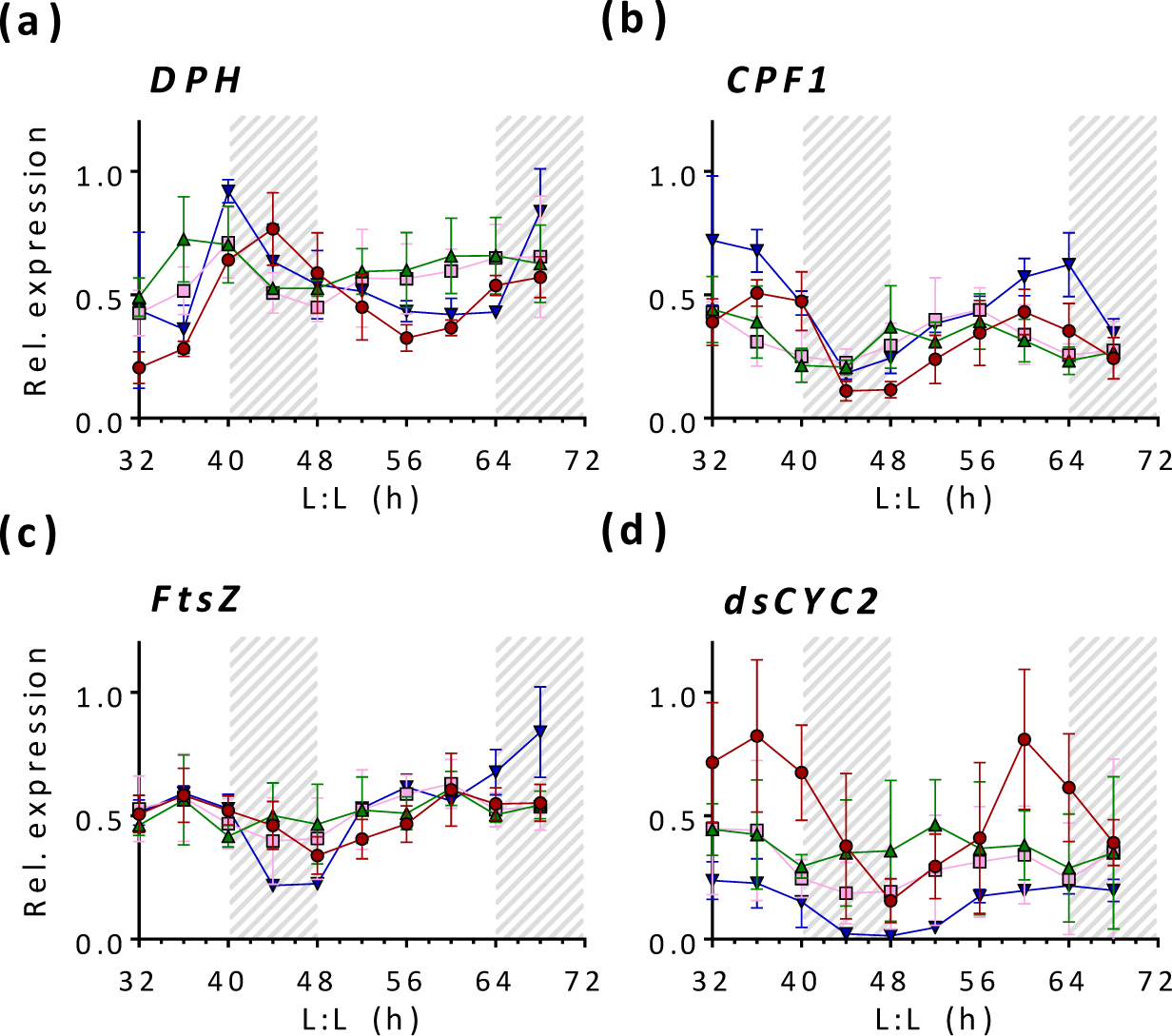
Analysis of the rhythmic expression of selected genes by qRT-PCR in *P. tricornutum* WT, *RITMO1* OE1, KO2 and KO2-C2 in free running conditions. Expression profiles of the photoreceptors *DPH* (a) and *CPF1* (b), the *FtsZ* (c) and the cyclin ds*CYC2* (d) for WT, KO2, OE1 and KO2-C2 lines. Expression values represent the average of three biological triplicates ±SD and are normalized using the *RPS* and *TBP* reference genes. Expression values are given relative to the maximum expression for each gene, where ‘1’ represents the highest expression value of the time series. Dots represent mean values, error bars represent SD (n=3). Grey dashed regions represent subjective nights in free-run conditions.

**Fig. S13:**
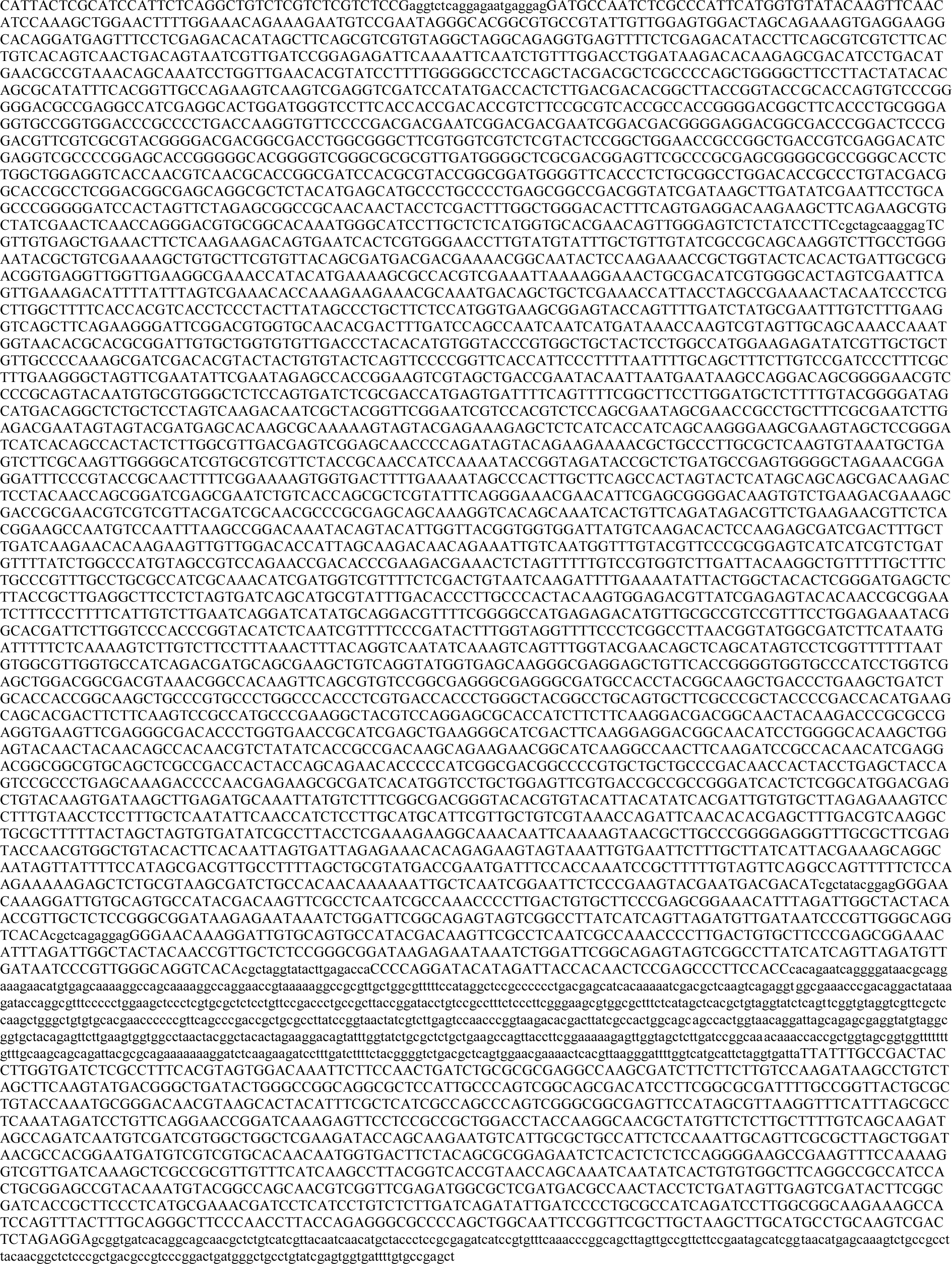
Sequence of the pL2-1 plasmid used to transform *RITMO1* KO2 cells to generate the complemented KO2-C2 strain.

**Fig. S14:**
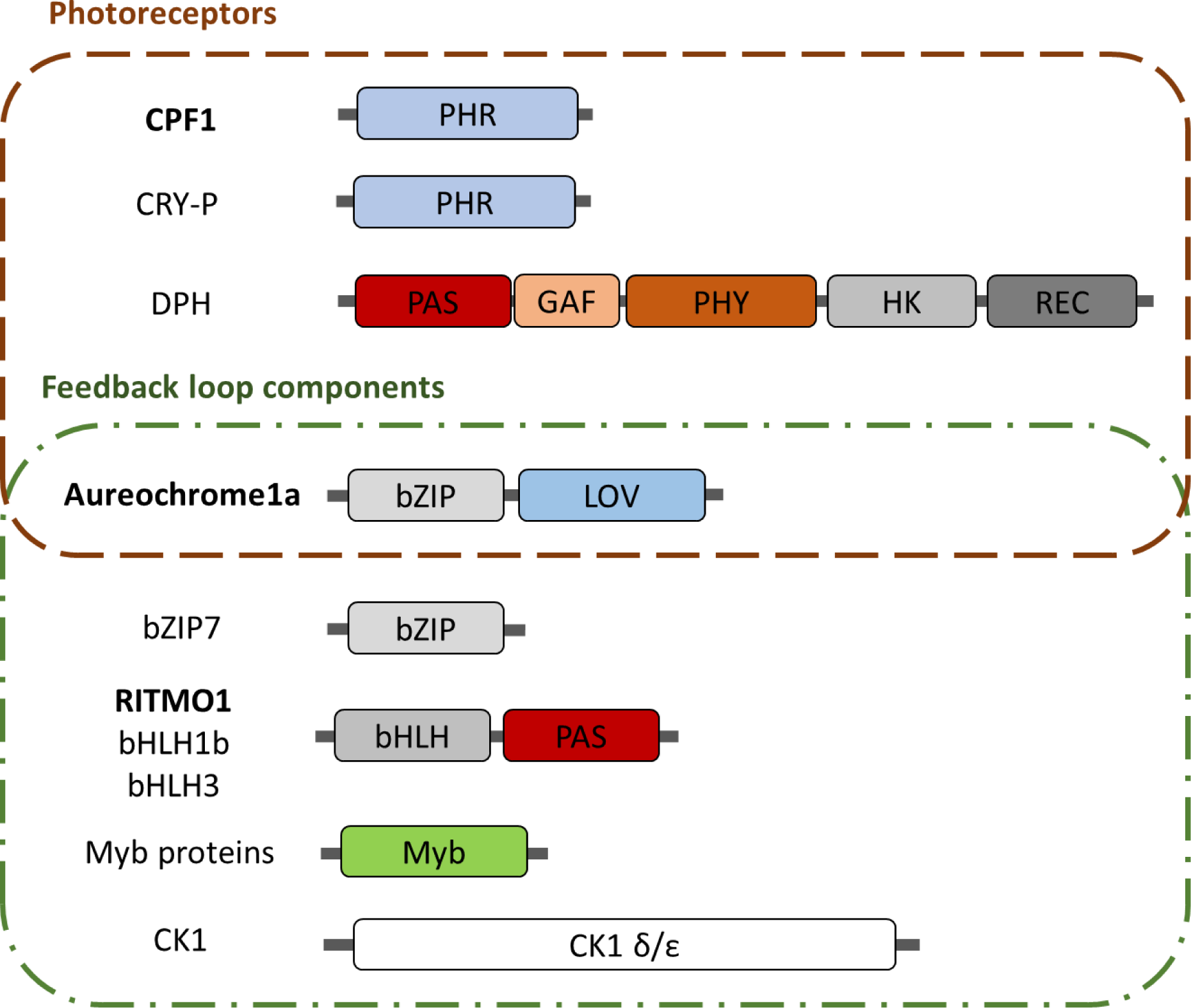
Scheme of the validated and putative components of the diatom circadian clock system. The figure includes diatom photoreceptors, putative component of the input pathway, and factors of the clock regulatory loop, identified in this study, Coesel *et al*., 2009; Annunziata *et al*., 2019; Madhuri *et al*., 2024 and diatom genomic investigations. The validated clock components are indicated in bold. The involvement of CPF1 in circadian clock has so far been shown only in heterologous mammalian cell system (Coesel *et al*., 2009). From top to bottom, proteins with photoreceptor domains CPF1 (Phatr3_J27429), Cry-P (Phatr3_J54342), DPH (Phatr3_J54330), the Aureochrome1a with both bZIP transcription factor (TF) and LOV photoreceptor domains (Phatr3_J8113), TFs containing bZIP domain bZIP7 (Phatr3_J48800), the animal-like bHLH-PAS domains in RITMO1 (Phatr3_J44962), bHLH1b (Phatr3_J44963, and bHLH3 (Phatr3_J42586); untested genes containing SANT/Myb domain, like the plant clock components. Diatom genomes also contain several proteins of the plant clock system with a CCT domain. However, these proteins miss other domains necessary for circadian clock function, and therefore are not included in the proposed scheme. CK1 δ/ε type like genes, known post-translational regulator of circadian clock across taxa, have been also found (Phatr3_J42322, Phatr3_J51110). The identifiers of the different components in the *P. tricornutum* genome are indicated (https://www.diatomicsbase.bio.ens.psl.eu/) (Villar *et al*., 2024).

